# A transient CD87-centred axis enhances the Th17 potential of cDC2 in neonates

**DOI:** 10.64898/2026.06.12.731823

**Authors:** Ramin Shakiba, Doğuş Altunöz, Kaushikk Ravi Rengarajan, Steven X. Cho, Yuxuan Zhang, Mert Meydanci, Josephine C. Owen, Hamsa Narasimhan, Nikos E. Papaioannou, Aikaterini Symeonidi, Silke Oberknapp, Sabine Schwamberger, Raymond Kiu, Lindsay J. Hall, Maria Colomé-Tatché, Dirk Haller, Jan P. Böttcher, Claudia A. Nold-Petry, Anne B. Krug, Marcel F. Nold, Barbara U. Schraml

## Abstract

Type 2 conventional dendritic cells (cDC2) orchestrate T cell immunity, yet the signals that regulate their tissue-specific functions across development remain poorly defined. We demonstrate that splenic cDC2/DC3 subset heterogeneity is established perinatally and that splenic ESAM^hi^ cDC2A undergo a transcriptional and functional transition around the time of weaning, which occurs independently of microbiota. Comparative transcriptomics identified neonatal-specific regulatory programs, including elevated expression of *Plaur*, encoding CD87/uPAR. CD87 sensitizes neonatal ESAM^hi^ cDC2A to coagulation factor XII (FXII), thereby enhancing their capacity to promote Th17 differentiation. In human infants, CD87 expression was high on cDC2, its expression declined with age and in plasma of preterm infants CD87 positively correlated with Th17-associated cytokines. Together, we identify a conserved link between coagulation pathways and cDC2 developmental programming that may offer new opportunities to modulate Th17 responses in infancy.

**Summary:** Splenic cDC2 undergo a transcriptional and functional remodeling around weaning. Elevated CD87 expression on neonatal cDC2 enhances their FXII-driven Th17 potential. Conserved in human infants, elevated CD87 expression reveals a coagulation-linked pathway shaping early-life dendritic cell function and suggests targets to improve vaccination.

## Introduction

The neonatal period is characterized by immune tolerance, yet newborns are capable of mounting protective immune responses against pathogens and after vaccination (Hong & Medzhitov, 2023; Kollmann et al., 2017). Type 2 conventional dendritic cells (cDC2) are potent coordinators of immunity that in neonates exhibit distinct functional characteristics, including distinct epitope bias, cytokine production and expression of co-stimulatory molecules, characteristics that are thought to facilitate tolerance to commensal microbes and environmental antigens (Hornef & Torow, 2020; Levy, 2007; Papaioannou et al., 2018). Accumulating evidence, however, indicates that rather than being intrinsically immature, neonatal cDC2 are functionally programmed by their tissue environment - signals including commensals, cytokines, and other developmental cues shape their phenotype in a tissue - and age-dependent manner (Bachus et al., 2019; de Kleer et al., 2016; Gollwitzer et al., 2014; Papaioannou et al., 2021; Torow et al., 2023; Weckel et al., 2023; Xing et al., 2026). Highlighting the existence of tightly regulated, context-specific instructive programs in early life, cDC2 immunocompetency emerges asynchronously across tissues and developmental stages (de Kleer et al., 2016; Papaioannou et al., 2021; Torow et al., 2023; Weckel et al., 2023; Xing et al., 2026).

Commensal encounter and dietary changes during the transition from breastfeeding to solid food at weaning critically impact immune development (Al Nabhani et al., 2019; Al Nabhani & Eberl, 2020; Hornef & Torow, 2020). While commensal- and diet-induced factors shape cDC function across tissues (Altunöz et al., 2026 *Preprint*; Gollwitzer et al., 2014; Köhler et al., 2021; Nguyen et al., 2025; Schaupp et al., 2020; Smout et al., 2023), less is known about how systemic host-derived factors contribute to the developmental programming of cDC2 function. The spleen is the primary site of blood filtration, allowing immune cells to monitor blood for the presence of pathogens and abnormal cells (Lewis et al., 2019). The strategic positioning of cDC2 in blood-exposed regions around the white pulp augments their ability to capture antigens and subsequently drive T helper (Th) responses upon migration to the T cell areas (Calabro et al., 2016; Liu et al., 2020; Yi & Cyster, 2013). This localization also positions cDC2 in niches that may allow them to sense and respond to circulating host-derived mediators generated during tissue remodeling, vascular activation, or coagulation.

Splenic cDC2 uniformly express CD11b and CD172a but can be divided into Notch2-dependent ESAM^hi^ or T-bet-expressing cDC2A, and Notch2-independent ESAM^low^ or T-bet-negative cDC2B (Baber et al., 2025; Brown et al., 2019; Hoelting et al., 2025; Lewis et al., 2011; Minutti et al., 2024). Both subtypes can induce Th17 responses, but ESAM^hi^ cDC2A more potently induce Tfh responses, while some evidence suggest ESAM^low^ cDC2B may better promote Th2 responses (Baber et al., 2025; Briseño et al., 2018; Hoelting et al., 2025; Lewis et al., 2011; Tussiwand et al., 2015). Recently, monocytic cDC2B-like cells have been described and it has been suggested to name these cells DC3 (Liu et al., 2023), although it remains unclear they are a functionally distinct lineage or a cell state (Amon et al., 2025; Hoelting et al., 2025). Notably, analogous subsets are conserved in humans, where CD1c^+^ DCs in peripheral blood comprise both cDC2 and DC3 populations (Dutertre et al., 2019; Villani et al., 2017).

In adult mice ESAM^hi^ cDC2A and ESAM^low^ cDC2B arise predominately from committed CLEC9A-expressing committed myeloid-derived bone marrow progenitors but they can also arise from transitional DCs (tDCs) with lymphoid features (Minutti et al., 2024; Papaioannou et al., 2021; Rodrigues et al., 2024; Schraml et al., 2013; Sulczewski et al., 2023). A putative lymphoid contribution to ESAM^hi^ cDC2A and ESAM^low^ cDC2B is prominent in the perinatal spleen (Papaioannou et al., 2021). Developmentally distinct cDC2 arising from myeloid and lymphoid progenitors in early life are transcriptionally and functionally similar (Papaioannou et al., 2021). Compared to cDC2 in adult life, however, cDC2 from neonatal mice are transcriptionally distinct and better at supporting T helper (Th) 17 and T regulatory (Treg) cell differentiation (Papaioannou et al., 2021). While both ESAM^hi^ cDC2A and ESAM^low^ cDC2B display heightened Th17 potential in early life, increased Treg potential was specifically observed in ESAM^hi^ cDC2A (Papaioannou et al., 2021). Age-dependent differences in gene expression are at least in part caused by IFN-I (Papaioannou et al., 2021) but the specific signals regulating splenic cDC2 function across age, as well as the turning point when cDC2 function changes during development are unclear.

Here, we show that splenic cDC2 undergo a transcriptional and functional switch around weaning that is regulated independent of microbiota. Comparative transcriptomics identified neonatal-specific regulators, including elevated expression of *Plaur* (encoding for CD87, also known as uPAR - urokinase plasminogen activator receptor), a receptor linking coagulation factor XII (FXII) to cAMP signaling and Th17-polarizing cytokine production (Göbel et al., 2016). CD87 expression was selectively increased in ESAM^hi^ cDC2A in neonatal compared to adult spleen, declined with age, and functionally sensitized ESAM^hi^ cDC2A to FXII, thereby enhancing their capacity to drive Th17 differentiation. Importantly, this axis is conserved in humans, with higher CD87 expression on CD1c^+^ DCs in infants compared to adults. Together, our findings uncover a previously unappreciated link between hemostatic pathways and the functional programming of antigen presenting cells in early life.

## Results

### Neonatal cDC2 undergo a functional switch around weaning

Having shown that cDC2 from neonatal mice are better at supporting Th17 differentiation than their adult counterparts (Papaioannou et al., 2021), we first wondered when this switch in cDC2 functionality takes place. We thus compared the ability of CD11b^+^ cDC2 (including both ESAM^hi^ cDC2A and ESAM^low^ cDC2B) sorted from mice at 2, 4 and 10 weeks post-birth to stimulate naïve OT-II cells (Supp. Fig. 1A, B). In all experiments pups were separated from their mothers at exactly three weeks of age. cDC2 from all ages were pulsed with Ovalbumin peptide 323-339 (OVA_323-339_) for 3 hours and then cultured with CellTrace Violet (CTV)-labelled OT-II transgenic T cells under Th17 polarizing conditions (Supp. Fig. 1C). Three days later OT-II cell proliferation and cytokine production were analyzed. CD11b^+^ cDC2 from all ages induced similar proliferation of OT-II cells, as measured by the number of dividing cells, as well as proliferation and division indices (Fig. 1A, Supp. Fig. 1D). As expected, cDC2 from 2-week-old mice induced more IL-17A production from OT-II cells than cDC2 from adult mice (Papaioannou et al., 2021) and the same was observed for IL-17F production (Fig. 1B). Interestingly, increased IL-17A and IL-17F production was also observed when OT-II cells were cultured with cDC2 from 2-week-old compared to 4-week-old mice (Fig. 1B). In contrast, TNF production from OT-II cells was similar when cultured with cDC2 from different ages (Fig. 1B). Similar results were obtained when ESAM^hi^ cDC2A from mice at 2, 4 and 10 weeks were sorted and cultured with OT-II cells as above (Supp. Fig. 1B, E). These data confirm that cDC2 from 2-week-old mice induce similar T cell proliferation and TNF production from T cells as their adult counterparts but exhibit qualitative differences in supporting Th17 differentiation. By 4 weeks of age splenic cDC2 have undergone a functional switch and exhibit adult Th17 potential.

**Figure 1:**
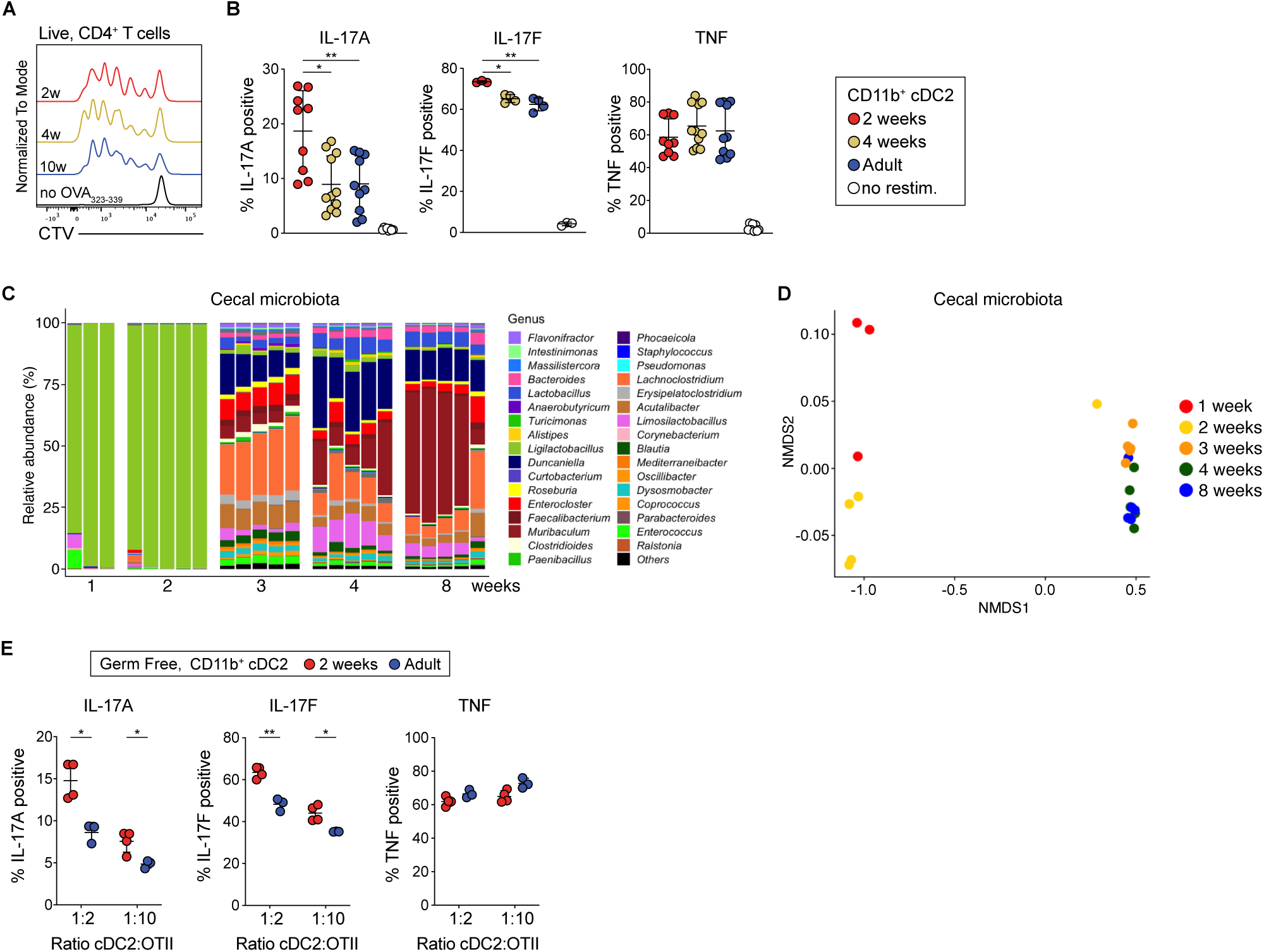
Splenic cDC2 undergo a functional switch around weaning. (A-B) Sorted splenic CD11c^+^MHCII^+^CD11b^+^ cDC2 from mice of indicated ages were pulsed with OVA_323-339_ and co-cultured with CTV-labelled OT-II cells under Th17 polarizing conditions at a 1:2 ratio (cDC2:OT-II). 3.5 days later OT-II cells were analyzed for cytokine production and CTV. **(A)** Representative CTV traces. **(B)** The percentage of IL-17A, IL-17F and TNF-positive cells within CTV^neg^ OT-II cells is shown. **(C)** Relative abundance of cecal microbiota from mice of indicated ages profiled via shotgun sequencing on the genus level. Bars represent biological replicates. **(D)** Non-metric multidimensional scaling (NMDS) analysis of microbiome community structure at genus level. (**E)** Splenic CD11b^+^ cDC2 from 2-week-old and adult germ-free mice were co-cultured with CTV-labelled OT-II cells under Th17 polarizing conditions at the indicated DC2:OT-II ratios as in **(A)**. The percentage of IL-17A, IL-17F and TNF-positive cells within CTV^neg^ OT-II cells is shown. Each dot represents one biological replicate. Horizontal bars represent mean, error bars represent SD. Data for IL-17A and TNF in **(B)** is pooled from three independent experiments. Data for IL-17F is pooled from two independent experiments. Data from **(E)** is from one independent experiment. Only statistically significant comparisons are indicated. Statistical analysis for (**B)** was performed using one-way ANOVA, for (**E)** was performed using two-tailed Welch’s t tests with correction for multiple comparisons. *p<0.05, **p<0.01, ***p<0.001, ****p<0.0001.

Mice typically start eating chow from around two weeks of age (Bailoo et al., 2020). The timing of the observed functional switch therefore correlates with weaning, which demarcates the transition from milk to grain-based chow diet in mice. Profiling cecal contents of wild type mice at 1, 2, 3, 4 and 8 weeks old using metagenomics showed an increase in microbial diversity across age, with the most prominent changes being evident between 2 and 3 weeks of age (Fig. 1C, D). These data confirm that weaning correlates with a strong increase in microbial diversity (Al Nabhani et al., 2019; van Best et al., 2020). Dietary changes at weaning, as well as presence of commensals have been linked to increased serum or tissue IFN⍺ and IFNγ levels in humans and mice (Al Nabhani et al., 2019; Erttmann et al., 2022; Goehring et al., 2016; Schaupp et al., 2020; Vatanen et al., 2022). Since cDC2 from adult compared to young mice show enrichment of IFN-regulated genes and *Ifnar1*-deficiency phenotypically rejuvenates splenic cDC2 from adult mice (Papaioannou et al., 2021), we hypothesized that IFNs suppress the Th17 potential of splenic cDC2 post weaning. To address this, we sorted splenic cDC2 from adult *Ifnar1^-/-^* mice and *CD11c^Cre^Stat1^flox/flox^* mice, which lack responsiveness to IFN-I or all IFNs, respectively, and cultured them with OT-II cells under Th17 polarizing conditions as described above. We observed no differences in the ability of cDC2 with impaired IFN responsiveness to induce T cell proliferation and IL17A/F or TNF production compared to cDC2 from control mice (Supp. Fig. 1F-H). Thus, loss of IFN signaling does not rejuvenate the Th17 potential of cDC2, indicating that weaning-associated changes in IFNs do not cause the switch in Th17 potential. Additionally, we isolated cDC2 from 2-week-old and adult germ-free (GF) mice and assessed their ability to induce T cell proliferation and Th17 effector differentiation. cDC2 from 2-week-old and adult GF mice induced similar OT-II proliferation and TNF production from OT-II cells under Th17 polarizing conditions (Fig. 1E, Supp. Fig. 1I). However, cDC2 from 2-week-old GF mice induced stronger IL-17A and IL17F production from OT-II cells than their adult counterparts (Fig. 1E). Thus, differences in Th17 potential between neonatal and adult cDC2 are programmed independently of commensal microbiota. Taken together, these data suggest that rather than the adult microenvironment suppressing Th17 potential of cDC2, the increased Th17 potential may be imprinted in the neonatal period by environmental or host-derived factors specifically present in early life.

### Splenic cDC2 transcriptionally change around weaning

To understand the transcriptional regulation of cDC2 around weaning we analyzed cDC2 and tDCs from a single cell RNA-sequencing (scRNA-seq) dataset of CD11c^+^MHCII^+^ splenocytes isolated from 1-, 3-, 4- and 6-week-old *Clec9a^Cre^Rosa^TOM^* myeloid DC-tracer mice (Fig. 2A) (Altunöz et al., 2026 *Preprint*). In these mice both *TOMATO* fluorescence signal and transcript levels allow tracing of cDCs originating from *Clec9a*-expressing myeloid cDC progenitors (Papaioannou et al., 2021; Schraml et al., 2013). All mice were weaned at exactly three weeks of age (Fig. 2A). cDC2 from all time points clustered together, indicating that variation was not driven by batch effects (Fig. 2B). Clusters 6 and 7 transcriptionally resembled ESAM^hi^ cDC2A and cluster 3 resembled tDCs (Fig. 2B, C, Supp. Fig. 2A, B, Supp. Table 1). Three clusters transcriptionally resembled ESAM^low^ cDC2B (cluster 2, 5 and 9), of which clusters 2 and 5 also showed similarity to DC3 (Fig. 2C, D, Supp. Table 1). ESAM^low^ cDC2B and DC3 reportedly resemble each other and an inability to clearly distinguish DC3 and cDC2B could be augmented in neonatal mice because DC3 and cDC2A/B gene signatures were generated by profiling cells from adult mice (Liu et al., 2023; Nguyen et al., 2025; Rodrigues et al., 2023). To further establish relatedness of neonatal cDC2 to cDC2A, cDC2B and DC3, we integrated our scRNA-seq dataset with 5 publicly available splenic cDC data sets (Brown et al., 2019; Liu et al., 2023; Narasimhan et al., 2025; Rodrigues et al., 2024; Sulczewski et al., 2023). Original cell annotations were used, except for Sulzcewski et al., which we annotated according to gene signatures provided in the original annotation (Supp. Fig. 3A-D). Visualization of the integrated data demonstrated that cells annotated as the same broad cell types in the individual datasets (cDC1, cDC2A, CCR7^+^ DCs, tDCs and pDCs) localized together in UMAP space (Supp. Fig. 4A). Contribution to Leiden clusters was driven by cell type rather than dataset or sequencing method, confirming accurate integration of cells with similar transcriptional properties (Supp. Fig. 4B-D). Cells of cluster 7 ESAM^hi^ cDC2A (7_DC2_ESAM^hi^_age) were found in Leiden cluster 3, together with DC2 from Liu et al., cDC2A from Rodrigues et al. and “cluster 14 mixed” of Brown et al., that contained mostly Tbet^+^ cDC2A but also some Tbet^neg^ cDC2. Cells of cluster 6 ESAM^hi^ cDC2A (6_DC2_ESAM^hi^_age) were equally divided between Leiden clusters 2 and 4 together with cells annotated as cDC2A from Brown et al., Rodrigues et al. and Narasimhan et al. These data confirm clusters 6 and 7 from our current analyses correspond to cDC2A (Supp. Fig. 4D). Cells of cluster 2 (2_DC2_ESAM^low^_age) were found in Leiden cluster 1, which also contained cDC2B from Brown et al. and cells annotated at CX3CR1^+^ cDC2B/DC3 by Rodrigues et al. Few cells of cluster 5 (5_DC2_ESAM^low^_age) were found in Leiden cluster 1, while most cells were found in Leiden cluster 18, together with DC3 from Liu et al. Interestingly, Leiden cluster 18 also contained a fraction of cells annotated as CX3CR1^+^ cDC2B/DC3 by Rodrigues et al., in line with the observation that fate mapping is required to accurately distinguish cDC2B from DC3 (Nguyen et al., 2025; Rodrigues et al., 2024). Cells of cluster 9 (9_DC2_ESAM^low^_age) were mostly found in Leiden cluster 12, together with cDC2B from Brown et al. and cDC2_4 annotated as cDC2B in Narasimhan et al. Taken together, this integration analysis confirmed that cluster 6 and 7 constitute ESAM^hi^ cDC2A, while cluster 5 most likely corresponds to DC3 and clusters 2 and 9 mostly resemble cDC2B (Supp. Fig. 4D).

**Figure 2:**
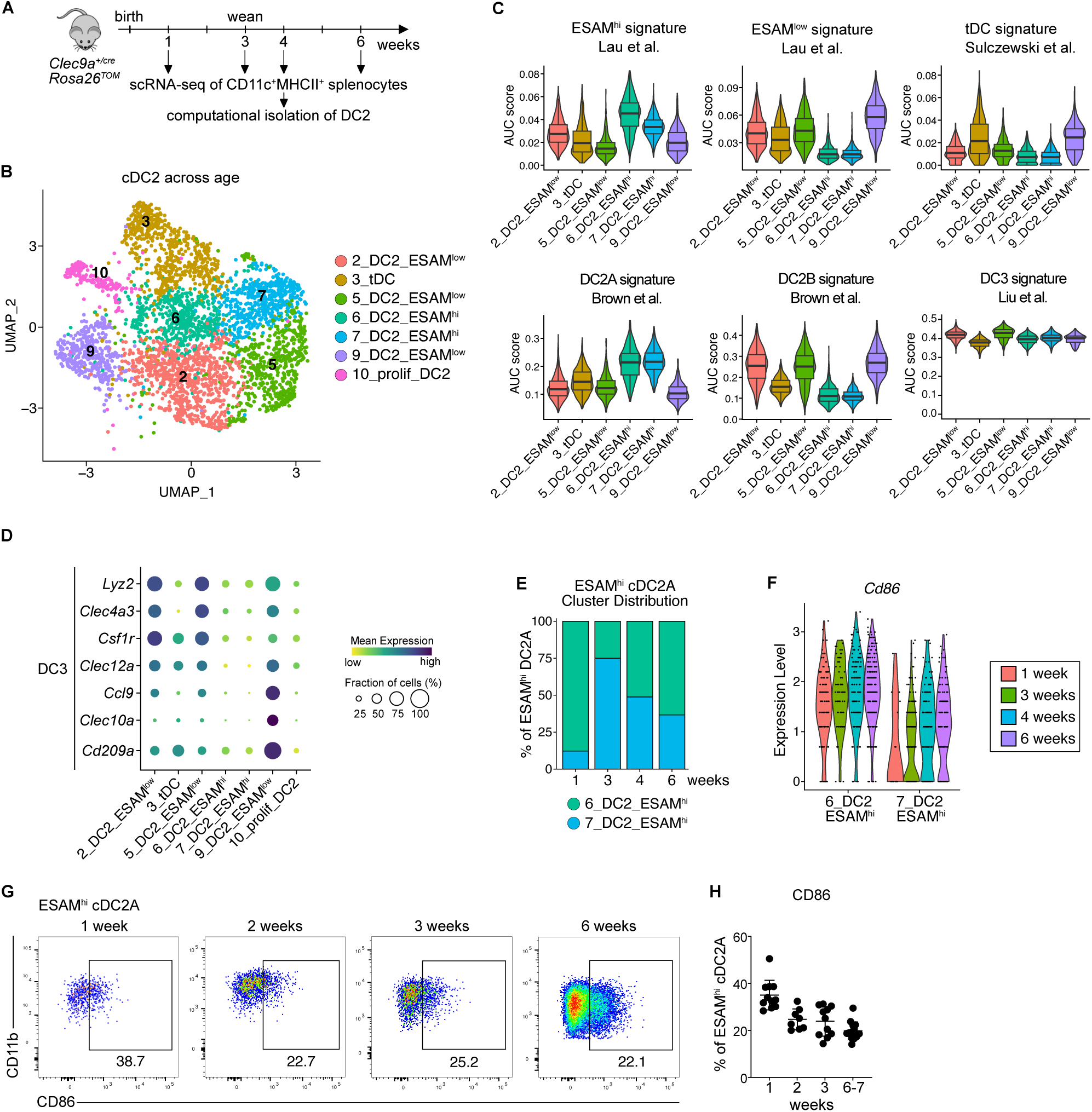
Weaning correlates with transcriptional changes in spleen cDC2. **(A)** cDC2 and tDC were computationally isolated from a scRNA-seq data set (Altunöz et al., 2026 *Preprint*). **(B)** UMAP display of 3716 cDC2 and tDCs grouped from all ages and annotated by cell type. **(C)** AUC scores for indicated signatures across cDC2 clusters. Box represents interquartile range, horizontal bar represents median, whiskers represent minimum and maximum enrichment scores. **(D)** Bubble plot displaying expression of indicated genes across clusters. **(E)** Relative frequency of ESAM^hi^ cDC2 clusters 6 and 7 at the different time points. **(F)** Expression of *Cd86* across times points in ESAM^hi^ cDC2 clusters 6 and 7. **(G-H)** Flow cytometry **(G)** and quantification **(H)** of the percentage of CD86 expressing ESAM^hi^ cDC2A. Data in **(H)** are pooled from 6 independent experiments. Each dot represents one biological replicate. Horizontal bars represent mean, error bars represent SD.

**Figure 3:**
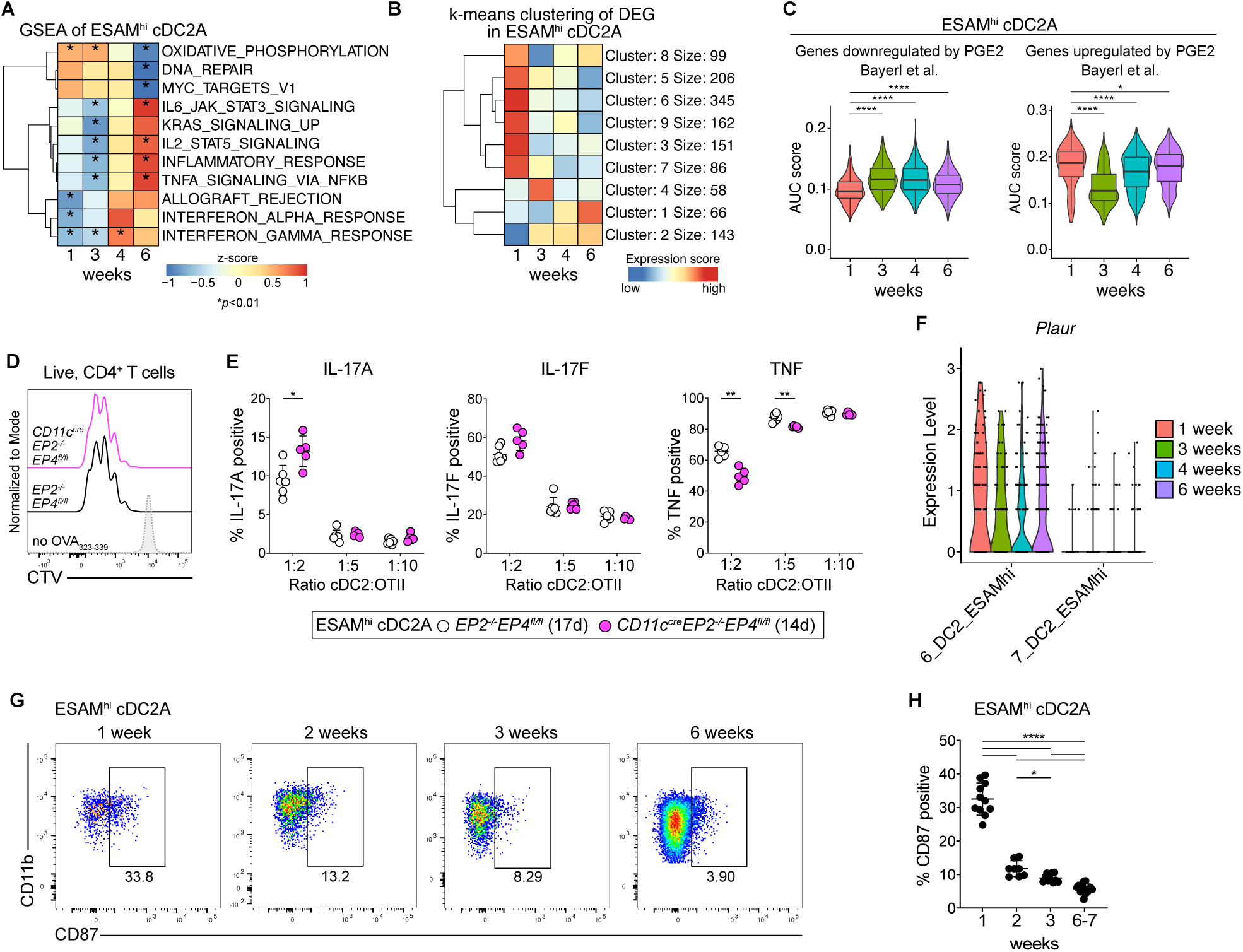
Identification of putative neonatal-specific regulators of cDC2 function. (A, **B)** GSEA of ESAM^hi^ cDC2 clusters 6 and 7 across age. Only significantly enriched hallmark gene sets are shown. **(B)** Genes differentially expressed in ESAM^hi^ cDC2 (clusters 6 and 7) between time points were analyzed. K-means clustering of these genes is shown. **(C)** AUC scores for genes regulated in cDCs by PGE2 signaling (Bayerl et al., 2023). Boxes represent interquartile range, horizontal bars represent median, whiskers represent minimum and maximum. **(D-E)** ESAM^hi^ cDC2A from mice of indicated genotypes were pulsed with OVA_323-339_ and co-cultured with CTV-labelled OT-II cells under Th17 polarizing conditions at the indicated DC:APC ratios for 3.5 days. **(D)** Representative CTV traces from one sample of the 1:2 ratio. **(E)** Percentage of IL-17A, IL-17F and TNF-positive cells within CTV^neg^ OT-II cells. Data is representative of three independent experiments. **(F)** Expression of *Plaur* across ESAM^hi^ cDC2 clusters 6 and 7 across time. **(G-H)** Representative gating **(G)** and relative frequency **(H)** of CD87 expression on ESAM^hi^ cDC2A at the indicated ages. Each dot represents one biological replicate pooled from 6 independent experiments. Horizontal bars represent mean, error bars represent SD. Statistical analysis was performed using multiple t-tests corrected for multiple comparisons using the Holm-Šídák method **(C)**, two-tailed Welch’s t tests with correction for multiple comparisons **(E)** and using one-way ANOVA **(H)**. Only statistically significant comparisons are indicated. *p<0.05, **p<0.01, ***p<0.001, ****p<0.0001.

**Figure 4:**
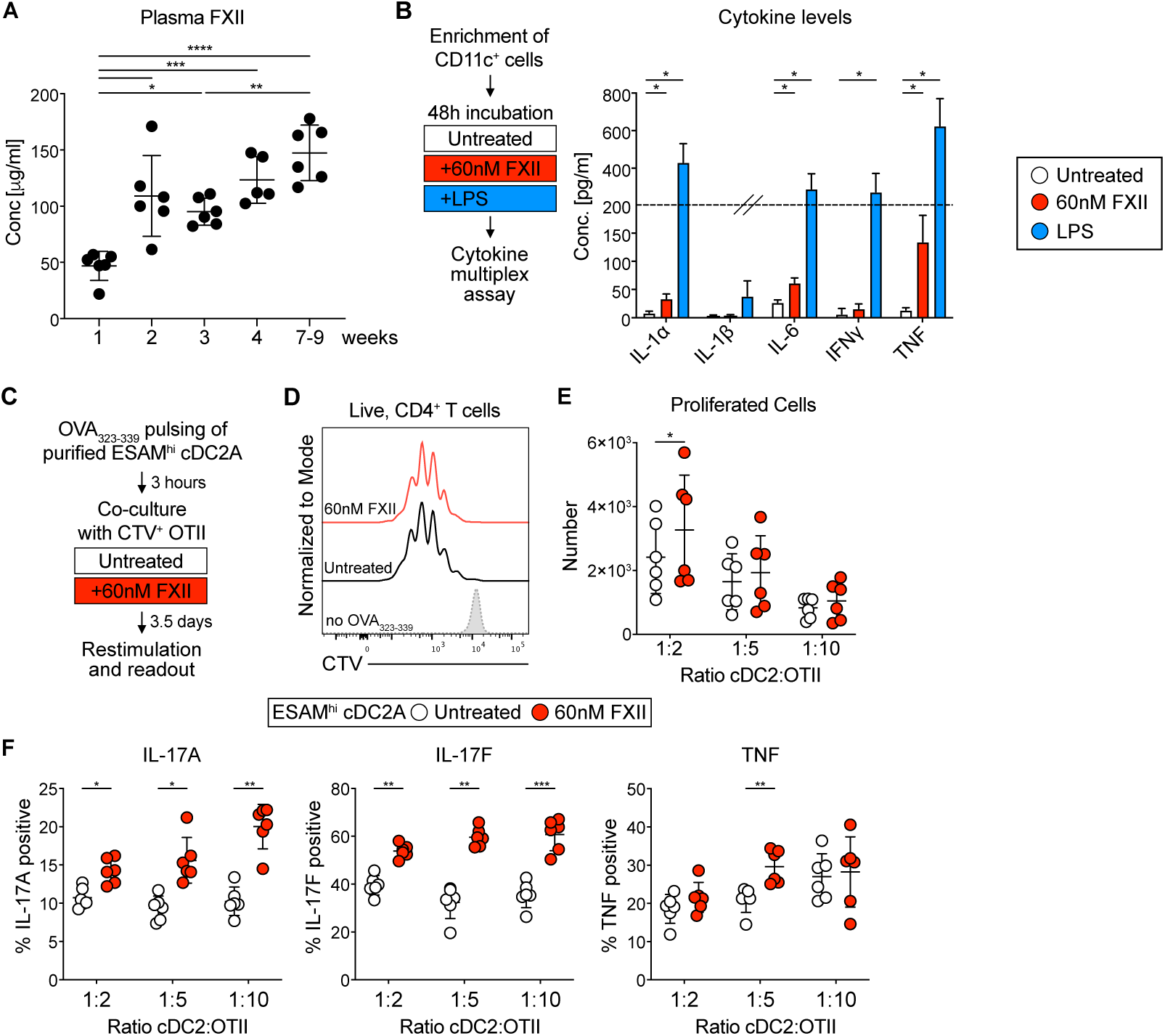
FXII treatment enhances the Th17 potential of neonatal cDC2. **(A)** FXII plasma concentration in mice of the indicated ages. **(B)** CD11c-enriched splenocytes from adult mice (16 weeks old) were left untreated or cultured with FXII or LPS for 48 hours. Cytokines were measured in supernatants by bead array (n=3). **(C-F)** Splenic ESAM^hi^ cDC2A from 2-week-old mice were pulsed with OVA_323-339_ and co-cultured with CTV-labelled OT-II cells under Th17 polarizing conditions at different DC:T cell ratios with or without 60nM FXII. 3.5 days later T cell cytokine production and CTV signal were analyzed. **(C)** Experimental scheme. **(D)** Representative CTV traces from the 1:2 ratio. **(E)** Number of CTV^neg^ OT-II cells. **(F)** Percentage of IL-17A, IL-17F and TNF-positive cells within CTV^neg^ OT-II cells. Each dot represents one biological replicate. Horizontal bars represent mean, error bars represent SD. Data in **B-F** are representative of two independent experiments. Statistical analysis: one-way ANOVA **(A)**, multiple paired t test **(B),** and two-tailed Welch’s t tests with correction for multiple comparisons **(E, F).** Only statistically significant comparisons are indicated. *p<0.05, **p<0.01, ***p<0.001, ****p<0.0001.

cDC2A have been defined as T-bet reporter-positive cDC2 but these cells are also largely ESAM positive (Brown et al., 2019; Minutti et al., 2024). Splenic cDC2 can be delineated into ESAM^hi^ and ESAM^low^ subsets by flow cytometry directly from birth (Lewis et al., 2011; Papaioannou et al., 2021). In the absence of a T-bet reporter, we thus adopt the ESAM^hi^ cDC2A and ESAM^low^ cDC2B nomenclature for the remainder of this manuscript, acknowledging that ESAM^low^ cDC2 may constitute a mixture of cDC2B and DC3. Because of this heterogeneity, we focused subsequent analyses on ESAM^hi^ cDC2A, which can be clearly delineated transcriptionally and by flow cytometry across age and show heightened Th17 potential in neonates (Supp. Fig. 1E) (Papaioannou et al., 2021).

Within ESAM^hi^ cDC2A the relative abundance of clusters 6 and 7 changed with age, with the relative abundance of cluster 6 being highest at week 1 (Fig. 2E). Cluster 6 expressed higher levels of CD86 than cluster 7 (Fig. 2F), suggesting it represents a distinct activation state of ESAM^hi^ cDC2A (Fig. 2F). Indeed, flow cytometry confirmed expression of CD86 on a fraction of ESAM^high^ cDC2A and the frequency of CD86-expressing ESAM^hi^ cDC2A was highest at week 1 compared to later time points (Fig. 2G, H, Supp. Fig. 2E), validating the suitability of the scRNA-seq approach to identify age-dependent differences in gene expression. In *Clec9a^Cre^Rosa^TOM^* mice *TOMATO* expression traces the fate of *Clec9a*-expressing cDC committed progenitors (Salei et al., 2020; Schraml et al., 2013). By computing the ratio of predicted transcripts of the unrecombined Rosa26 locus and *TOMATO* reads, we confirmed that TOMATO expression in all cDC2 clusters was low at week one and increased with age (Supp. Fig. 2C, D). By three weeks of age, most cDC2 were *TOMATO*-positive as expected (Papaioannou et al., 2021) (Supp. Fig. 2C, D). Comparison of TOM^neg^ and TOM^+^ cells within individual clusters was not possible at the 1-week timepoint as cell numbers were limiting and at the other time points the unrecombined *Rosa26* locus was identified as a commonly differentially expressed gene (Supp. Table 2). Few other genes, if any, were identified as differentially expressed between TOM^neg^ and TOM^+^ cells across clusters (Supp. Table 2), confirming our previous results showing that ontogeny does not strongly contribute to transcriptional heterogeneity and that developmentally distinct cDC2 in early life are transcriptionally similar (Papaioannou et al., 2021). Accordingly, we pooled the TOM^neg^ and TOM^+^ ESAM^hi^ cDC2A for subsequent age-dependent transcriptional comparisons, thereby increasing statistical power through a larger combined cell count.

### Identification of putative neonatal-specific regulators of cDC2 function

To gain insights into specific transcriptional pathways that could regulate ESAM^hi^ cDC2A with age, we first performed gene set enrichment analyses (GSEA). GSEA identified several pathways enriched in ESAM^hi^ cDC2A (clusters 6 and 7 combined) from 4 and 6-week-old mice compared to earlier time points, including genes downstream of IFNα and IFNγ (Fig. 3A). Pathways enriched at 1-week-old compared to other time points, included oxidative phosphorylation, DNA repair and MYC targets (Fig. 3A). Bulk-sequencing of cDC2 from adult compared to 2-week-old mice had equally demonstrated regulation of these pathways (Papaioannou et al., 2021) and our data show that ESAM^hi^ cDC2A shows these transcriptional changes as early as around weaning.

Because the above results did not clearly point us to neonatal-specific regulators of ESAM^hi^ cDC2A function, we performed comparative gene expression analysis to identify genes differentially expressed across age. We combined ESAM^hi^ cDC2A clusters 6 and 7 and performing pairwise comparisons from each time point against all other time points. K-means clustering of these genes revealed 9 clusters with distinct expression pattern across age (Fig. 3B, Supp. Table 3). Most clusters contained genes with highest expression at one week of age (clusters 3, 5, 6, 7, 8, 9), while genes in cluster 4 showed highest expression at 3 weeks of age and expression of genes in clusters 1 and 2 increased with age (Fig. 3B). Cluster 8 contained *Crem* (cyclic AMP-responsive element modulator) that mediates cAMP signaling, which in cDC2 is activated in response to prostaglandin E2 (PGE2) or coagulation factor XII (FXII) and promotes Th17 potential (Göbel et al., 2016; Lee et al., 2020).

Interestingly, ESAM^hi^ cDC2A from 1-week-old mice exhibited increased regulation of PGE2-responsive genes (Bayerl et al., 2023) (Fig. 3C), leading us to address whether constitutive PGE2 signaling in cDC2 may underlie the increased Th17 potential of cDC2 in neonates. We sorted splenic ESAM^hi^ cDC2A from 2-week-old *Itgax^Cre^Ptger2^-/-^Ptger4^fl/fl^* mice, which have a deletion of EP4 in CD11c-expressing cDCs on a global EP2-deficient background, thus conditionally eliminating PGE2 signaling in cDCs. Although the transcriptional data was obtained from 1-week-old mice, we used 2-week-old mice to obtain sufficient cells for functional assays and because we had previously observed heightened Th17 potential of ESAM^hi^ cDC2A at this age. *Ptger2^-/-^Ptger4^fl/fl^* mice were used as control. ESAM^hi^ cDC2A from both genotypes were co-cultured with OT-II cells under Th17 polarizing conditions for three days. Interestingly, cDC2 from both genotypes induced comparable OT-II cell proliferation, whereas IL17A production from OT-II cells was higher when cultured with ESAM^hi^ cDC2A from *Ptger2^-/-^Ptger4^fl/fl^*compared to control mice (Fig. 3D, E, Supp. Fig. 5A). These data indicate that constitutive PGE2 signaling does not increase the Th17 potential of neonatal cDC2.

**Figure 5:**
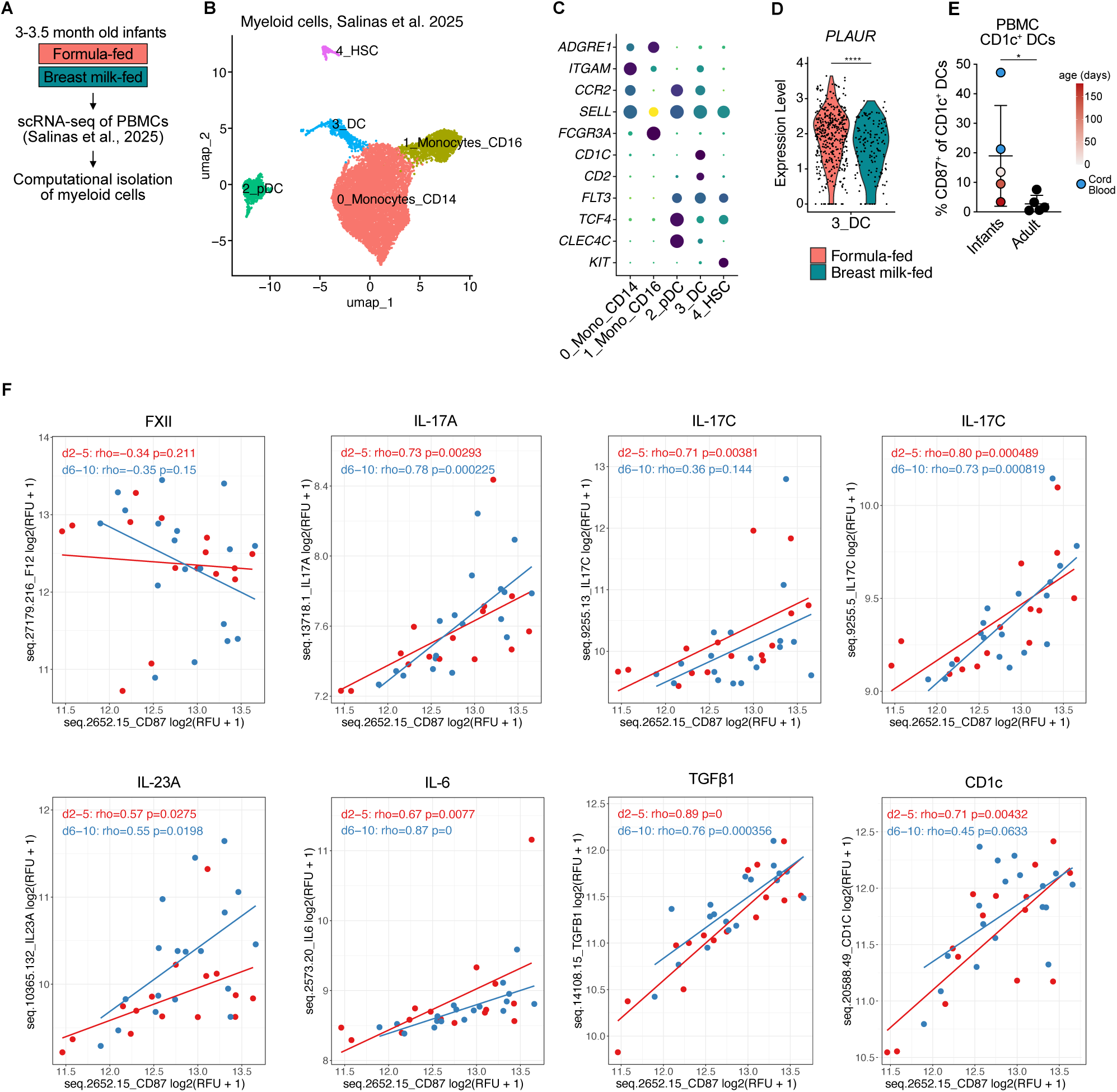
CD87 expression is increased in cDC2 from infants compared to adults. (A-D) scRNA-seq data of PBMCs of infants (Salinas et al., 2025) was analyzed. **(A)** Scheme of computational isolation of myeloid cells from infant PBMCs. **(B)** UMAP display of computationally isolated myeloid cells annotated using cell type specific genes displayed in **(C). (D)** Expression of PLAUR in the CD1c^+^ DC cluster stratified by diet. **(E)** PBMCs from infants (age stratified by color) and adults were profiled by flow cytometry. The relative frequency of CD87 expressing cells within CD1c^+^ DCs was quantified. **(F)** Preterm plasma was collected and assayed by proteomics before being stratified into two timepoints, early (day of life 2–5, n =15) and late (day of life 6–10, n =18). Spearman correlation analysis was performed between CD87 and indicated proteins. RFU = relative fluorescence units. Statistical analysis for **D, E** was performed using Student’s t-test. *p<0.05, **p<0.01, ***p<0.001, ****p<0.0001.

Coagulation FXII, also known as Hageman factor, induces cAMP signaling and promotes Th17 potential in cDCs by engaging the glycosylphosphatidylinositol (GPI)-anchored CD87 (also known as urokinase-type plasminogen activator receptor – uPAR) (Göbel et al., 2016). CD87 is encoded by *Plaur* and was found in k-means gene cluster 3 with highest expression at week 1 (Supp. Table 3). *Plaur* expression was higher in cluster 6_ESAM^hi^ cDC2A, which was the dominant cDC2A cluster at 1 week of age, compared to cluster 7_ESAM^hi^ cDC2A (Fig. 2E, Fig. 3F). Flow cytometry confirmed these transcriptional data with CD87 expression being highest on ESAM^hi^ cDC2A from 1-week-old mice and declining thereafter (Fig. 3G, H, Supp. Fig. 5B), suggesting that the FXII-CD87 axis might be involved in regulating the Th17 potential of ESAM^hi^ cDC2A from neonates.

### FXII treatment enhances the Th17 potential of neonatal cDC2

FXII is a plasma protein that is produced as an inactive zymogen in the liver and gets activated to FXIIa upon contact with negatively charged surfaces (Long et al., 2016). Both the active FXIIa and zymogen FXII induce cAMP signaling in CD11c^+^ cDCs from adult mice in a CD87-dependent manner (Göbel et al., 2016). We detected FXII in plasma of neonatal mice, indicating that ESAM^hi^ cDC2A could potentially be exposed to FXII in neonates (Fig. 4A). FXII concentrations gradually increased postnatally (Fig. 4A), in line with the known gradual postnatal development of the coagulation system (LaRusch et al., 2010; Monagle & Massicotte, 2011; Schmidt et al., 1998).

FXII treatment stimulates production of the Th17 promoting cytokines IL-6 and IL-23 from human peripheral blood mononuclear cells (PBMCs) and boosts production of these cytokines from lipopolysaccharide (LPS)-treated murine CD11c^+^ cDCs (Göbel et al., 2016). To assess whether FXII treatment alone is sufficient to induce cytokine production from murine cDCs, we cultured CD11c-enriched splenocytes with or without FXII for 48 hours as described (Göbel et al., 2016) and measured cytokines in supernatants by bead array (Fig. 4B). Because detection of cytokines in supernatants requires sufficient input cells, CD11c-enriched splenocytes from adult mice were used. LPS served as positive control, inducing production of TNF, IL-6, IFNγ, and IL-1α as expected. FXII treatment alone also induced production of IL-6, IL-1α and TNF when compared to unstimulated control cells (Fig. 4B), confirming that FXII is biologically active in the zymogen form and induces cytokine production from CD11c-enriched cells *in vitro*.

To directly assess whether FXII can modulate the function of neonatal ESAM^hi^ cDC2A, we performed T cell co-cultures. We sorted ESAM^hi^ cDC2A from 2-week-old mice and cultured them with OT-II cells at different cDC:T cell ratios under Th17 conditions as above, either in the presence or absence of FXII (Fig. 4C). Addition of FXII to co-cultures did not influence the proliferation of T cells (Fig. 4D, E, Supp. Fig. 6A). However, at each cDC:T cell ratio we investigated, we observed increased IL-17A and IL-17F production from OT-II cells in the presence of FXII compared to in the absence of FXII, while TNF production was similar in the presence or absence of FXII (Fig. 4F). Thus, addition of FXII to neonatal ESAM^hi^ cDC2A does not quantitatively alter their T cell activation potential but specifically boosts the ability of ESAM^hi^ cDC2A to induce Th17 differentiation (Fig. 4F). These findings identify the CD87–FXII axis as a putative regulator of ESAM^hi^ cDC2A function in neonates.

### CD87 expression is increased in CD1c^+^ cDCs from infants compared to adults

We next assessed if CD87 could be relevant in the functional regulation of cDC2 in human infants. To address whether infant cDC2 express *PLAUR* (encoding CD87, aka PLAUR), we computationally isolated myeloid cells from a published scRNA-seq dataset of PBMCs from 3-3.5 months old infants (Salinas et al., 2025) (Fig. 5A, B). Based on the expression of *FLT3*, *CD1C* and *CD2,* we were able to distinguish CD1c^+^ cDCs (Fig. 5B, C), which comprise both cDC2 and DC3 (Dutertre et al., 2019; Villani et al., 2017). Because this data set contained cells from infants that were either exclusively breast fed or formula fed, we also stratified the *CD1C^+^*cDC cluster by diet. *PLAUR* was found on CD1c^+^ cDCs in both conditions, although transcript levels were higher in infants on formula versus breast milk (Fig. 5D). To further validate these findings and assess whether CD87 expression on human CD1c^+^ cDCs is regulated with age, we profiled PBMCs of 5 healthy adult donors and 5 infants (gestational age between 26-36 weeks, sampled from cord blood to 6 months postnatal age) by flow cytometry. Indeed, we found higher frequency of CD87 expressing CD1c^+^ cDCs in infants compared to adults and that CD87 frequency decreased with postnatal age (Fig. 5E, Supp. Fig. 7A).

Preterm infants are at high risk of bleeding and infections delays in adjusting to extrauterine life in the hemostatic and immune systems, yet, they can mount Th17 responses and some studies even found increased circulating Th17 cells in preterm compared to term infants (Khizroeva et al., 2023; Lao et al., 2022; Melville & Moss, 2013; Rito et al., 2017). Because CD87 can be shed from the cell surface into the circulation (Blasi & Carmeliet, 2002; Mahmood et al., 2018), we investigated whether plasma CD87 is correlated with the abundance of Th17 proteins in preterm infants. We performed targeted aptamer-based proteomic assay on plasma from preterm infants (n=22, gestational age 28^+2^ weeks [IQR 25^+5^ – 32^+2^]) from two postnatal timepoints – days 2-5 of life and days 6-10 of life – designated as early and late timepoints, respectively. We detected CD87 in all samples. Targeted correlation analyses segregated by time point revealed that plasma abundance of CD87 positively correlated with CD1c at the early time point. At both time points, CD87 positively correlated with the Th17- inducing cytokines IL-6, TGFβ and IL-23A, as well as with the Th17 effector cytokines IL-17A and IL-17C (Fig. 5F). No correlation was observed for FXII (Fig. 5F). Thus, we observe a positive correlation between CD87, CD1c and Th17-associated cytokines, all together indicating that the age-related changes in CD87 abundance on cDC may have functional relevance, such as shaping cDC functions.

## Discussion

Although neonates are at increased risk of infection, they exhibit an intrinsic propensity towards Th17 responses. Here, using comparative transcriptomic analyses we reveal neonatal-specific transcriptional features in ESAM^hi^ cDC2A, including elevated expression of CD87, a receptor that links coagulation FXII to the enhanced Th17 potential of neonatal ESAM^hi^ cDC2A. Notably, this pathway appears conserved in humans, as cDC2 from infants exhibit higher CD87 expression compared to adults and plasma CD87 positively correlates with the cDC2 marker CD1c and Th17-associated cytokines in plasma of preterm infants. Collectively, these findings identify a previously unrecognized connection between hemostatic signaling pathways and the developmental programming of cDC2 function in early life that contributes to a neonatal Th17 bias and might prove relevant to improve infant survival and vaccination strategies.

While it has been recognized that splenic cDC2 from young mice exhibit heightened Th17 potential (Papaioannou et al., 2021), we here show that these cells undergo transcriptional and functional transition towards an adult-like Th17 potential around the time of weaning. This functional switch occurred independently of microbiota, despite the well-established influence of weaning-associated dietary and commensal changes on the functions of immune cells (Al Nabhani et al., 2019; Altunöz et al., 2026 *Preprint*; Gollwitzer et al., 2014; Schaupp et al., 2020; van Best et al., 2020). In line with prior work showing higher IFN-regulated genes in cDC2 from adult compared to young mice (Papaioannou et al., 2021), scRNA-sequencing demonstrated enrichment of IFN-regulated genes in ESAM^hi^ cDC2A already from 4 weeks post-birth. Although *Ifnar1*-deficiency phenotypically rejuvenates splenic cDC2 from adult mice (Papaioannou et al., 2021), we found that loss of STAT1, and thus IFN-responsiveness, did not rejuvenate the Th17 potential of cDC2 from adult mice. These observations suggested that the Th17 potential of cDC2 might not be suppressed by postweaning IFN signaling, but rather could be actively imprinted by the neonatal environment.

scRNA-seq of cDCs across age identified 5 distinct clusters of cDC2-like cells. Using published gene signatures generated in adult mice (Brown et al., 2019; Lau et al., 2016), we could identify 2 clusters of ESAM^hi^ cDC2A already present in neonatal spleen. Distinction of the remaining 3 clusters into cDC2B and DC3 was not possible using published gene signatures alone but was facilitated by integration with other transcriptional datasets containing splenic cDC subsets from adult and neonatal mice (Brown et al., 2019; Liu et al., 2023; Narasimhan et al., 2025; Rodrigues et al., 2024; Sulczewski et al., 2023). Through this analysis we identified 2 clusters resembling cDC2B and one cluster resembling DC3 (Baber et al., 2025; Liu et al., 2023), altogether suggesting that the full heterogeneity of splenic cDC2/DC3 subsets is present from birth. Interestingly, all cDC2/DC3 clusters showed low frequency of *Clec9a^Cre^*-driven *TOMATO* transcript at one week-old compared to later time points, suggesting that DC3, like cDC2 may exhibit layered ontogeny. Transcriptional comparison showed that *TOMATO^+^* cDC2/DC3 arising from committed myeloid cDC progenitors are largely indistinguishable from *TOMATO^neg^* cells within the same clusters, supporting the notion that distinct progenitors converge onto a common terminal cDC2 state distinguishable only by fate mapping (Baber et al., 2025; Rodrigues et al., 2024; Sulczewski et al., 2023). Such ontogenetic conversion highlights the necessity to further define the environmental signals that determine cDC2 fate and function in tissues, considering that various subset-defining markers are responsive to cytokine signals (Amon et al., 2025; Brown et al., 2019; Nguyen et al., 2025).

Because ESAM^hi^ cDC2A can be identified in the spleen by flow cytometry from birth, we focused age-dependent transcriptional comparisons on these cells. Interestingly, we identified *Crem* as a gene selectively increased in ESAM^hi^ cDC2A from 1-week-old mice compared to other time points. CREM mediates cAMP signaling-induced transcriptional changes and cAMP signaling downstream of PGE2 and CD87 to boost the Th17 potential of cDC2 (Göbel et al., 2016; Lee et al., 2020). CD87 is a pleiotropic GPI-anchored membrane protein that binds urokinase plasminogen activator (uPA) and FXII and by pairing with other proteins, including integrins, regulates various cellular functions, including cell migration, proliferation, survival, and invasion (Stoppelli, 2013). In line with the known function of CD87 in promoting the Th17 potential of cDCs in autoimmune encephalomyelitis in mice (Göbel et al., 2016), we found that FXII specifically boosted the potential of neonatal cDC2 to induce IL-17A and IL-17F production from OT-II cells, while not affecting their ability to drive T cell proliferation and TNF production. In our system FXII was added throughout the co-cultures and could thus conceivably act directly on T cells, which also express CD87. However, it is reported that FXII does not influence the differentiation and polarization of CD4^+^ T cells (Göbel et al., 2016).

Suggesting conserved functional regulation of cDC2, we observed higher expression of CD87 in cDC2 from PBMCs of infants compared to adults and found plasma CD87 to positively correlate with CD1c and Th17-promoting cytokines in the plasma of preterm infants. Plasma CD87 is correlated with inflammation in the context of some viral infections and cancer (Blasi & Carmeliet, 2002; Mahmood et al., 2018) and in COVID-19 patients has been linked to an inflammation-associated population of myeloid cells (Sarif et al., 2021). A positive correlation between CD87 and CD1c in plasma of preterm infants thus suggests CD1c⁺ DCs as a potential source of CD87. Although CD1c is reported to exist as alternatively spliced and secreted isoform (Woolfson & Milstein, 1994), its presence in serum could also reflect the rapid turnover of neonatal cDCs or release from cell debris. Following vaccination, *PLAUR* expression is upregulated in PB-monocytes of infants (Nouri et al., 2023). Together with our observation that FXII stimulation of CD11c-enriched cDCs induces IL-6 and TNF production, these findings suggest that the FXII–CD87 axis could be leveraged to enhance Th17 differentiation, for example in the context of vaccination, or to block it to dampen Th17-associated inflammation.

CD87 expression is induced by inflammation and in response to pathogen associated patterns, such as LPS (Coleman et al., 2001; Florquin et al., 2001; Göbel et al., 2016; Xue et al., 1998). In mice, neonatal CD87 expression on cDC2 declined rapidly between one and two weeks of age and a rapid decline in CD87 frequency was also observed in our human cohort. However, increasing sample size will be necessary to determine when a switch in CD87 expression occurs in human infants and to identify the signals that regulate its expression on neonatal cDC2. Increased expression of PLAUR on CD1c^+^ cDC2 from infants on formula over breast milk suggest that external factors like diet could have an influence on CD87 expression, which requires further investigation.

The physiologic concentrations of most coagulation proteins are known to increase with gestational age and postnatally (Monagle & Massicotte, 2011), in line with our observation that plasma FXII levels in mice increase with age. Beyond its role in hemostasis, the coagulation system actively shapes inflammation and immunity, a concept captured by immune-hemostatic crosstalk, which has been extensively studied in the context of immunothrombosis and thromboinflammation (Engelmann & Massberg, 2013; Long et al., 2016; Palumbo, 2022). In this context, higher expression of CD87 on ESAM^hi^ cDC2A in early life may increase their sensitivity to FXII, potentially amplifying its immunological effects despite lower systemic availability. Preferential positioning of cDC2 in blood exposed regions of the neonatal spleen may further enable access of ESAM^hi^ cDC2A to FXII present in blood (Liu et al., 2020). In this context, it is important to mention that the structural organization into red and white pulp in mice takes place between 1 and 2 weeks after birth (Cheng et al., 2019; Lewis et al., 2019) and systemic analysis of cDC2 positioning during this time has not been performed. CD87 itself, by associating with integrins, serves as an adhesion molecule (Tarui et al., 2001) and may further promote the positioning of ESAM^hi^ cDC2A in specific tissue niches. FXII also plays a role in pathogen defense and limits pathogen spread by trapping bacteria within fibrin clots (Nickel et al., 2025). Although to our knowledge, a direct adjuvant effect of FXII on cDC2 during bacterial infection has not been demonstrated, it will be interesting to investigate in the future if elevated expression of CD87 on neonatal ESAM^hi^ cDC2A enables immune surveillance against pathogens. Additionally, CD87 supports conversion of plasminogen to plasmin, thereby promoting the activation of latent TGFβ (Godár et al., 1999; Stoppelli, 2013). While TGFβ is closely linked to Th17 biology, it also stimulates epithelial to mesenchymal transition (Mangan et al., 2006; Miettinen et al., 1994), suggesting that ESAM^hi^ cDC2A could contribute to tissue remodeling in early life.

Taken together, our findings position early life as a distinct immunological window during which cDC2 function is actively shaped by non-microbial environmental cues, including components of the hemostatic system. By identifying a transient FXII–CD87 axis as a developmental regulator of cDC2 function, our work expands the concept of immune–hemostatic crosstalk to the steady-state programming of antigen presenting cells. From a translational perspective, the heightened CD87 expression in infant cDC2 and its inducibility in inflammatory settings raise the possibility that this pathway could be leveraged to modulate vaccine responses or susceptibility to infection in early life. Future studies will need to define the upstream signals controlling CD87 expression, the spatial and temporal dynamics of FXII availability in lymphoid tissues, and whether targeting this axis can safely enhance protective immunity without exacerbating inflammatory pathology.

## MATERIALS AND METHODS

### Mice

Clec9a^tm2.1(icre)Crs^ (*Clec9a^Cre^*; Jackson Laboratory Stock No: 025523), Ifnar1^tm1Agt^ (*Ifnar^−/−^)* (MMRRC Stock No: 32045-JAX), Tg(Itgax-cre)1-1Reiz (*Itgax^cre^*) (Jackson Laboratory Stock No: 018967), Stat1^tm1.1Mmul^ (*Stat1^fl/fl^*), OT-II (B6.Cg-Tg(TcraTcrb)425Cbn/J, Jackson Laboratory Stock No: 004194) and C57BL/6JRccHsd mice were bred and maintained at the Biomedical Center, LMU Munich, in individually vented cages with a 12 h dark/light cycle. *Ptger2^-/-^Ptger4^fl/fl^* mice were bred and maintained at the Klinikum Rechts der Isar, Technical University of Munich (TUM). Germ-free animals were bred at the ZIEL institute for Food & Health TUM, Freising, Germany in isolators and germ-free status was routinely monitored. Food and water were provided ad libitum. Male and female littermates were used in this study unless otherwise stated. In all experiments mice were separated from their mothers at 21 days after birth and adult mice used in this study were 8–12 weeks old, unless otherwise specified. Mice were euthanized by cervical dislocation or if younger than 3 weeks by decapitation. All animal procedures were performed in accordance with national and institutional guidelines for animal welfare and approved by the respective authorities.

### Flow Cytometry

Cells were incubated in 50 µL of FACS buffer containing an Fc receptor–blocking antibody (anti-mouse CD16/32) for 10 minutes at 4 °C. Then 50 µL of a 2X antibody mastermix containing fluorochrome-conjugated antibodies against surface markers along with fixable viability dye eFluor™ 780 (Thermo Fisher Scientific) was added and incubated for 25 minutes at 4 °C. Subsequently, cells were washed twice and resuspended in fresh FACS buffer prior to flow cytometry. Intracellular cytokine staining was performed after surface staining using the Intracellular Fixation & Permeabilization Buffer Kit (Thermo Fisher Scientific), intranuclear staining was performed with the FOXP3 Transcription Factor Staining Kit (Thermo Fisher Scientific) according to the manufacturer’s protocol. For quantification CountBright™ counting beads (Thermo Fisher Scientific) were used as described previously (Kumaraswami et al., 2020). Intracellular cytokine staining was done after stimulation with phorbol 12-myristate 13-acetate (PMA, 10 ng/mL, Calbiochem) and ionomycin (1 µg/mL, Sigma-Aldrich) at 37 °C for a total of 5 hours, with Brefeldin A (5 µg/mL, Biolegend) added during the final 3 hours of culture. Antibodies for flow cytometry are listed in Supp. Table 4. Samples were acquired on an LSR Fortessa cytometer (BD Biosciences) using BD FACSDiva software (version 8, BD Biosciences) and analyzed using FlowJo (Tree Star, Inc.). cDC2 were sorted on a FACSAria Fusion (BD Biosciences).

### Cell isolation

Spleens were cut into small pieces and enzymatically digested in 1 mL RPMI containing 200 U/mL Collagenase IV (Worthington) and 0.2 mg/mL DNase I (Roche) for 30 minutes at 37 °C with shaking (180 rpm). Suspensions were then passed through a 70 μm cell strainer, diluted with ice-cold FACS buffer (PBS 1% fetal calf serum [FCS], 2.5 mM EDTA, and 0.02% sodium azide) and centrifuged. Red blood cells were lysed for 2 min in ACK buffer (MilliQ water containing 1 mM EDTA, 1.55 M NH₄Cl, and 100 mM KHCO₃) at 4 °C. After washing with FACS buffer cells were resuspended in FACS buffer and filtered again through a 70 μm strainer before downstream analyses. For functional assays and scRNA-seq experiments, PBS supplemented with 1% FCS and 2.5 mM EDTA was used for cell isolation.

### In vitro stimulation with human FXII and cytokine quantification

Splenocytes were enzymatically digested and CD11c^+^ cells were enriched using CD11c MicroBeads (Miltenyi) according to the manufacturer’s protocol. 1 × 10⁵ cells were stimulated with 60 nM human FXII (Cell Systems) or 1 µg/mL LPS (*E. coli*, Enzo) in 100 µL complete RPMI (RPMI with 10% FCS, 1% penicillin/streptomycin, 1% non-essential amino acids, 1% sodium pyruvate, 1% L-glutamine, and 0.05 mM β-mercaptoethanol). After 48 hours cytokines were measured in the culture supernatants using the LEGENDplex™ Mouse Inflammation Panel (BioLegend). Only cytokines passing the limit of detection were quantified.

### DC:T-cell co-cultures

Total CD11b^+^ cDC2 or ESAM^hi^ cDC2A were sorted from CD11c^+^ enriched cells from spleens of mice at the indicated ages (Papaioannou et al., 2021) and pulsed for three hours in 96-well V-bottom plates with 10 µg/mL chicken OVA_323-339_ (InvivoGen) in complete RPMI. Cells were the washed twice to remove residual OVA_323-339_ and resuspended in complete RPMI medium, serially diluted, and co-cultured with 2500 naïve CellTrace™ Violet (CTV)-labelled OT-II at the indicated ratios. Naïve CTV-labeled OT-II cells were isolated from the spleens of adult mice with the MojoSort™ Mouse CD4 Naïve T Cell Isolation Kit (Biolegend) and labeled with CTV (Thermo Fisher Scientific) according to manufacturer’s instructions.

Cultures were supplemented with 5 ng/mL TGF-β, 20 ng/mL IL-6, 10 µg/mL anti-IL-4, and 10 µg/mL anti-IFN-γ for Th17 conditions (all Biolegend). For FXII stimulation experiments, the co-culture medium was supplemented with 60 nM human FXII (Cell Systems). After 3.5 days of culture cells were restimulated with 10 ng/mL PMA (Calbiochem) and 1 µg/mL ionomycin (Sigma-Aldrich) for 5 hours, Brefeldin A (5 µg/mL, Biolegend) was added for the last three hours. Cells were processed for flow cytometry as described above.

### FXII ELISA

Blood was collected from mice at different ages into plasma tubes containing lithium heparin (Li-heparin, 25 I.U./mL; Sarstedt) and gently mixed. Samples were centrifuged at 2000 x g for 10 minutes at room temperature, supernatants were transferred into 1.5-mL microcentrifuge tubes and centrifuged again to obtain cell-free plasma. Plasma was stored at -80 °C and diluted 1:250,000 for FXII quantification by mouse FXII ELISA kit (total FXII antigen; Abcam).

### Shotgun sequencing and taxonomic profiling of Cecal microbiome

The cecum was isolated, opened and cecal contents were transferred into Eppendorf tubes and snap frozen in liquid nitrogen. Samples were stored at -80°C until analysis. Briefly, after extraction of genomic DNA and sequencing, shotgun metagenome raw reads (FASTQ) were firstly trimmed, adaptor-removed and quality-filtered using fastp v0.20.0 (-q 20). Subsequently, host-associated sequences were removed via KneadData v0.10.0 with *mus musculus* genome (GRCm38) bowtie2 index file retrieved from https://benlangmead.github.io/aws-indexes/bowtie (with options --bypass-trim and –reorder) to generate clean FASTQ reads that consist of pure bacterial sequences. Next, Kraken v2.1.3 was utilized for taxonomic assignment for purified metagenome reads (Kraken2 standard RefSeq indexes retrieved from https://benlangmead.github.io/aws-indexes/k2, June 2022), with confidence level set at 0.1. Bracken v2.6.2 was then deployed to re-estimate the relative abundance of taxa at both genus level from Kraken2 outputs as recommended. To simplify data for visualization purpose, minimal genera with <1000 reads across all samples were classified as ‘Others’ to simplify representations prior to data visualization. Data visualization was performed in R using ggplot2 v3.5.2. NMDS plots were generated using vegan v2.7.2. MaAsLin2 (R library) was utilized to perform differential analysis at default parameters.

### Single cell RNA Sequencing analyses

cDC2 and tDC cluster were computationally isolated from a scRNAseq dataset of splenic CD11c^+^MHCII^+^ cells from 1-week, 3-week, 4-week, and 6-week-old *Clec9a^+/cre^Rosa^TOM^* mice (Altunöz et al., 2026 *Preprint*) and subjected to reanalysis using the *Seurat* (v4.3.0) package. The *sctransform* package was used to normalize, scale and find variable features of the dataset. Using the FindAllMarkers and FindMarkers commands of the Seurat package differentially expressed genes were identified through pairwise comparisons between combined ESAM^hi^ clusters of different ages. K-means clustering of the differentially expressed genes across age was performed with the pheatmap (version 1.0.13) package with nkmeans=9. AUCell (v.3.18) package was used to score cells for enrichment of published gene signatures (Supp. Table 1). Gene set enrichment analysis was performed using the singleseqgset package. Raw sequencing data of PBMCs from individual subjects from Salinas et al. (Salinas et al., 2025) was loaded separately into Seurat. Sequencing data from each subject was filtered to exclude genes present in less than 5 cells and to only include cells with less than 20% mitochondrial genes, more than 750 unique features, and 500 to 50,000 UMIs. Data from individual subjects was then normalized, and integrated using Harmony (Korsunsky et al., 2019), followed by dimensional reduction via UMAP. Myeloid cells were then identified based on specific genes provided in the original publication, isolated and subjected again to unsupervised clustering.

For dataset integration across studies, all datasets, along with the corresponding metadata, were downloaded from the respective repositories. scRNA-seq data from Sulczewski et al. were reanalyzed using Seurat with thresholds described in the original publication (Sulczewski et al., 2023). The *sctransform* package was used to normalize, scale and find variable features of the dataset. Dimensional reduction was performed via UMAP and Louvain clustering was used to cluster the data in an unsupervised manner. Annotation of unsupervised clusters were based on expression of marker genes provided in the original publication. All datasets were pre-processed with scanpy (Wolf et al., 2018), retaining only cells for which cell type annotation existed and using the same parameters; namely mitochondrial counts less than 20%, filtering cells with counts in less than 200 genes and removing genes that were found in less than 3 cells. The only exception was for the Rodrigues et al. dataset, whereas per the original publication cells with less than 1,000 genes and more than 7,000 genes were removed. In addition, B cells, T cells and NK cells were also removed.

For the integration of the different datasets a benchmark was performed, using the scIB package (Luecken et al., 2022), in order to identify the integration method that produces the most reliable results. We tested four different integration methods, namely Harmony (Korsunsky et al., 2019), scanorama (Hie et al., 2024), scVI (Gayoso et al., 2022) and drVI (Moinfar & Theis, 2026 *Preprint*). Based on the results of the benchmarking, drVI was the method that performed best in terms both of batch correction, but also in conservation of the biological signal. All integration methods were unaware of the cell type annotations. The integrated dataset (Supp. Fig. 4A) was further analyzed with scanpy. By extracting the 2,000 most variable genes, the genes were clustered using the Leiden algorithm within scanpy, with a resolution of 1.7. The clustering resulted in 38 clusters (Supp. Fig. 4B) and the respective cell types were projected on it (Supp. Fig. 4C).

The contribution of each dataset and each cell type annotation per cluster was visualized as a dot plot (Supp. Fig. 4D), where only for visualization purposes the cell types were hierarchically clustered using Ward linkage on Euclidean distances. Row and column ordering in visualizations was determined from dendrogram leaf order without applying discrete cluster partitioning.

### Human cohort

In order to examine the cellular expression of CD87 on neonatal and adult dendritic cells, blood samples were collected through the GLAM&I study GLAM&I study (The Gut, Lung and Their Microbiomes & Immunology; ethics approval RES-19-0000-883A). Cord blood was collected from 2 infants (GA = 26+2 & 29+0 weeks), peripheral blood was collected from 3 infants (GA = 26+5, 30+3 & 35+1weeks) at postnatal day of life 11, 135 & 179, and from 5 healthy adult volunteers. No active inflammation was detected at the time of sampling. For plasma proteomics a separate subset of infants, (n = 22; gestational age 28+2 weeks [IQR 25+5 – 32+2]) had blood collected at 2 postnatal timepoints, day of life 2-5 (n=15) and day of life 6-10 (n=18).

### Flow cytometry of human PBMCs

Immediately after collection, blood was centrifuged (300xg, 15min, room temperature) and plasma stored at -80°C. The remaining cells were diluted in 2ml of 1x DPBS (Dulbecco’s Phosphate-Buffered Saline, Life Technologies). Peripheral blood mononuclear cells (PBMCs) were isolated by layering diluted cells on warmed Histopaque-1077 (37°C) (Sigma-Aldrich) using a 15ml SepMate tube (Stemcell Technologies) before being centrifuged at room temperature at 1200xg for 10min. Collected PBMCs were then resuspended in 10ml of 1x DPBS and centrifuged at room temperature at 400xg for 15min. Supernatants were removed, and cell pellets resuspended in RPMI-1640 (Gibco) with 1% human serum (Equitech-Bio) and 1:500 Mycozap Plus-PR (Lonza). Samples were then centrifuged at 400xg for 15min at 4°C and washed again with 1ml of flow cytometry (FC) buffer (1x DPBS, 2% fetal bovine serum (FBS, Bovogen) and 2nM EDTA (Sigma-Aldrich)). Pellets were then resuspended to a total volume of 100μl. Then, 14μl from each sample was transferred into a separate 1.7ml microcentrifuge tube to create an unstained control. 9.5μl of human Fc-receptor binding inhibitor polyclonal antibody (Invitrogen) was added to each of the stained samples, and samples were incubated for 20min at 4°C in the dark. After incubating, samples were stained with the human surface-stain antibodies (Supp. Table 5) for 30min in the dark at 4°C, after which they were washed twice with 1ml of FC-buffer and samples stored in the dark at 4°C until acquisition.

Unstained and stained samples were acquired using the 5-Laser Cytek Aurora (Cytek Biosciences, Fremont, CA) flow cytometer configured with 5 lasers (355 nm, 405 nm, 488 nm, 561 nm, 640 nm) and 67 detectors (64 fluorescence channels, FSC, blue laser SSC, violet laser SSC). The Cytek Aurora flow cytometer was calibrated daily using SpectroFlo QC Beads (Cytek Biosciences). Samples were unmixed using SpectroFlo software (Cytek Biosciences). Data were analyzed using FlowJo software (v10.10.0, Ashland, OR, BD Life Science, USA). After removing all cell doublets and debris, live CD45^+^ cells were first classified into Myeloid antigen-presenting cells (APCs) as CD3^neg^CD19^neg^HLA-DR^+^ cells before being further gated to remove monocyte subsets and designating CD1c^+^ cDCs as CD3^neg^CD19^neg^HLA-DR^+^CD14^neg^CD16^neg^CD123^neg^CD11c^+^CD1c^+^. Further gating was then performed within CD1c^+^ cDCs to identify CD87^+^ cells (Supp. Fig. 7A).

### Preterm plasma proteomics

Immediately after blood collection, blood was centrifuged (300×g, 15min, room temperature) and plasma stored at -80°C until proteomic profiling using SomaScan 11K assay (v5.0, SomaLogic, Boulder, CO, USA), a semi-quantitative aptamer-based assay, was performed by SomaLogic. Aptamers called SOMAmers (Slow Off-rate Modified Aptamer) are single-stranded oligonucleotides designed for high affinity and specificity to protein targets. Briefly, SOMAmers immobilised on strepavidin-coated beads were incubated with plasma samples to capture target proteins before bound proteins were biotinylated and SOMAmer-protein complexes cleaved to be released into solution. Complexes were then recaptured and protein denatured to release SOMAmers for quantification via DNA hybridisation to complementary probes on microarrays to generate Cy3-relative fluorescence units (RFU), indicative of proportional abundance of protein. Samples were assayed at 3 serial dilutions (0.005%, 1%, 20%) depending on which target proteins were being assessed. Raw signal values were then processed and normalised for hybridisation efficacy, inter- and intra- plate variability using SomaLogic’s standard pipeline. Protein expression data were then log2-transformed prior to analysis. Spearman rank correlation was performed to assess associations between the PLAUR/CD87 (seq.2652.15) and proteins of interest, CD1C, FXII, IL-17A, IL-17C, IL-17F, TGF-β1 and IL-6. As some proteins had >1 aptamers, we selected only the aptamers where Somalogic has verified internally to bind target protein via pull-down SDS-PAGE or mass spectrometry (CD1c, seq.20588.49; FXII, seq.27179.216; IL-17A, seq.13178.1; IL-17C, seq.9255.13; IL-17C, seq.9255.5; TGF-β1, seq.14108.15) or where unavailable, we selected the aptamer with the most varied published cross-validation (IL-6, seq.2573.20; IL-23A, seq.10365.132). The exception to this selection was IL-17F where both aptamers (seq.14026.24, seq.2775.54) were missing both internal and external characterization. Analyses were conducted separately for early (day of life 2–5) and late (day of life 6–10) timepoints. Data analysis was performed in R version 4.4.0 (http://www.r-project.org).

### Statistical analysis

Statistical analyses were performed using Prism version 10 (GraphPad Software). Unless otherwise specified, comparisons between two groups were performed using a two-tailed unpaired t-test with Welch’s correction. Percentage data were logit-transformed prior to statistical testing. For multiple group comparisons, one-way analysis of variance (ANOVA) with Tukey’s multiple comparisons or multiple paired t test was performed. p-value of less than 0.05 was considered statistically significant. For comparing expression levels and AUCscores in scRNA-seq datasets statistical analysis was performed using multiple t tests corrected for multiple comparisons using the Holm-Šídák method.

## Acknowledgements

We thank members of the Schraml lab for helpful discussions and critical reading of the manuscript. We acknowledge the Core Facilities for Flow Cytometry, Bioimaging, Bioinformatics and Animal Models at the Biomedical Center, LMU Munich for providing equipment and expertise. This work has been funded by an ERC Starting Grant (ERC-2016-STG-715182) and by the Deutsche Forschungsgemeinschaft (DFG, German Research Foundation) - TRR 359—Project number 491676693 (project B05), FOR2599 (Project P03, SCHR 1444/2-1) and SCHR 1444/3-1. D.A. was supported by a YLSY Doctoral Scholarship from the Republic of Türkiye Ministry of National Education. C.A.N.P. was supported by NHMRC Investigator Leadership grant Application IDs 1173584 and 2033196. M.F.N. was supported by the Fielding Fellowship 2017 and by the Victorian Government’s Operational Infrastructure Support Program. The authors have no conflicting financial interests.

## AUTHOR CONTRIBUTION

R.S., D.A., K.R.R., H.N., S.X.C., Y.Z., M.M., N.E.P., S.O., and J.C.O. performed experiments and generated data. R.S., K.S. and M.C.T., performed bioinformatic analyses. R.K. and L.J.H. performed the taxonomic profiling of cecal microbiome analysis. J.C.O, S.X.C. and M.F.N. performed targeted plasma proteomics from preterm infants and analyzed the data. D.H, S.S., A.B.K., J.P.B. provided mice. S.X.C., M.C.T., M.N. and A.B.K. contributed new reagents/analytic tools/scientific input. R.S., D.A., and B.U.S. wrote the paper. B.U.S. conceptualized and supervised the study.

## DATA AND MATERIALS AVAILABILITY

All study data are included in the article and/or supporting information.

