## Supplemental information for "A transient CD87-centred axis enhances the Th17 potential of cDC2 in neonates"

### Supplementary Figure 1:

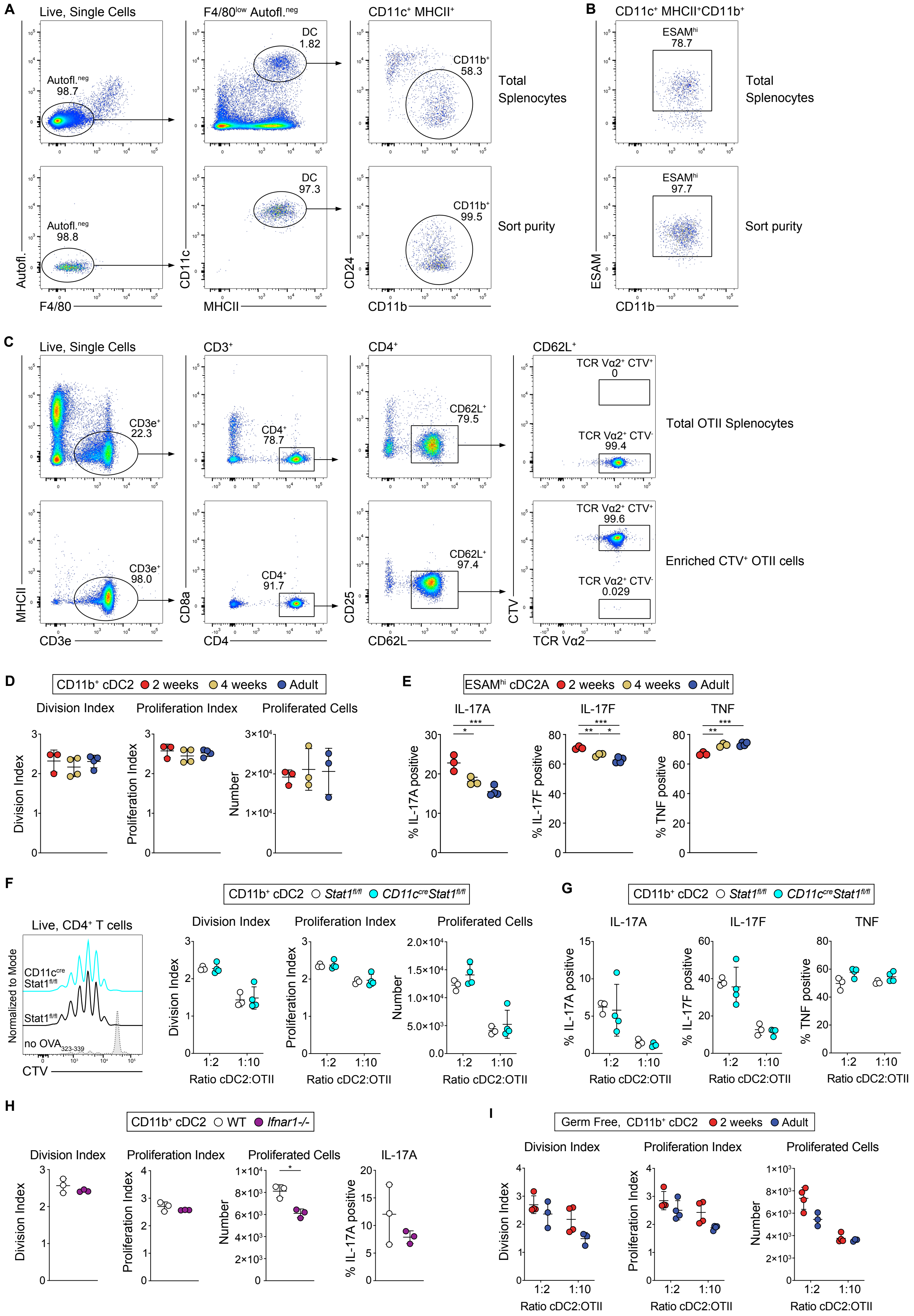

**Supplementary Figure 1:**

**(A, B)** Representative sort strategy and post-sort purity for $CD11b^{+}$ cDC2 **(A)** and $ESAM^{hi}$ cDC2A **(B)** from CD11c-enriched splenocytes showing results for an adult sample. Live leukocytes were gated and red pulp macrophages excluded by gating on $F4/80^{low}$ non-autofluorescent cells, from which cDCs were identified as $CD11c^{+}MHCII^{+}$ cells. Within cDCs, cDC2 were identified as $CD11b^{+}$ and if indicated accordingly further stratified by ESAM expression **(B)**. **(C)** Representative purity analysis of magnetically enriched OT-II cells from adult OT-II mice. From live leukocytes, T cells were gated as $CD3e^{+}$. From these, $CD4^{+}$ T cells with a $CD62L^{+}CD25^{neg}$ phenotype were identified as naïve and assessed for CTV labelling before co-culture. **(D)** Sorted splenic $CD11c^{+}MHCII^{+}CD11b^{+}$ cDC2 from mice of indicated ages were pulsed with $OVA_{323\text{-}339}$ and co-cultured with CTV-labelled OT-II cells under Th17 polarizing conditions at a 1:2 ratio (cDC2:OT-II). 3.5 days later the number of proliferated $CTV^{neg}$ cells) and division and proliferation indices were calculated. Data representative of two independent experiments. (**E)** Splenic $ESAM^{hi}$ cDC2A from mice of indicated ages were pulsed with $OVA_{323\text{-}339}$ and co-cultured with CTV-labelled OT-II cells under Th17 polarizing conditions at 1:2 ratio ($ESAM^{hi}$ cDC2A:OT-II). 3.5 days later percentage IL-17A, IL-17F and TNF-positive cells within $CTV^{neg}$ OT-II cells was calculated. **(F-I)** Sort-purified splenic $CD11c^{+}MHCII^{+}CD11b^{+}$ cDC2 from adult *Ifnar1*$^{-/-}$ mice **(H)**, *CD11c*$^{cre}$*Stat1*$^{fl/fl}$ mice **(F-G),** or germ-free (GF) mice of indicated ages **(I)** were pulsed with $OVA_{323\text{-}339}$ and co-cultured with CTV-labelled OT-II cells under Th17 polarizing conditions at the indicated cDC2:OT-II ratios. 3.5 days later, proliferated OT-II cells were restimulated with PMA and Ionomycin as above. Cell proliferation and IL-17A, IL-17F and TNF production from $CTV^{neg}$ OT-II cells was analyzed as above. Each dot represents one biological replicate. Data from **D,** is representative of three experiments. Data from **E, F, G** is representative of two independent experiments. Data from **H, I** is from one independent experiment. Horizontal bars represent mean, error bars represent SD. Statistical analysis was performed using two-tailed Welch's t tests with correction for multiple comparisons. *p<0.05, **p<0.01, ***p<0.001, ****p<0.0001.

### Supplementary Figure 2:

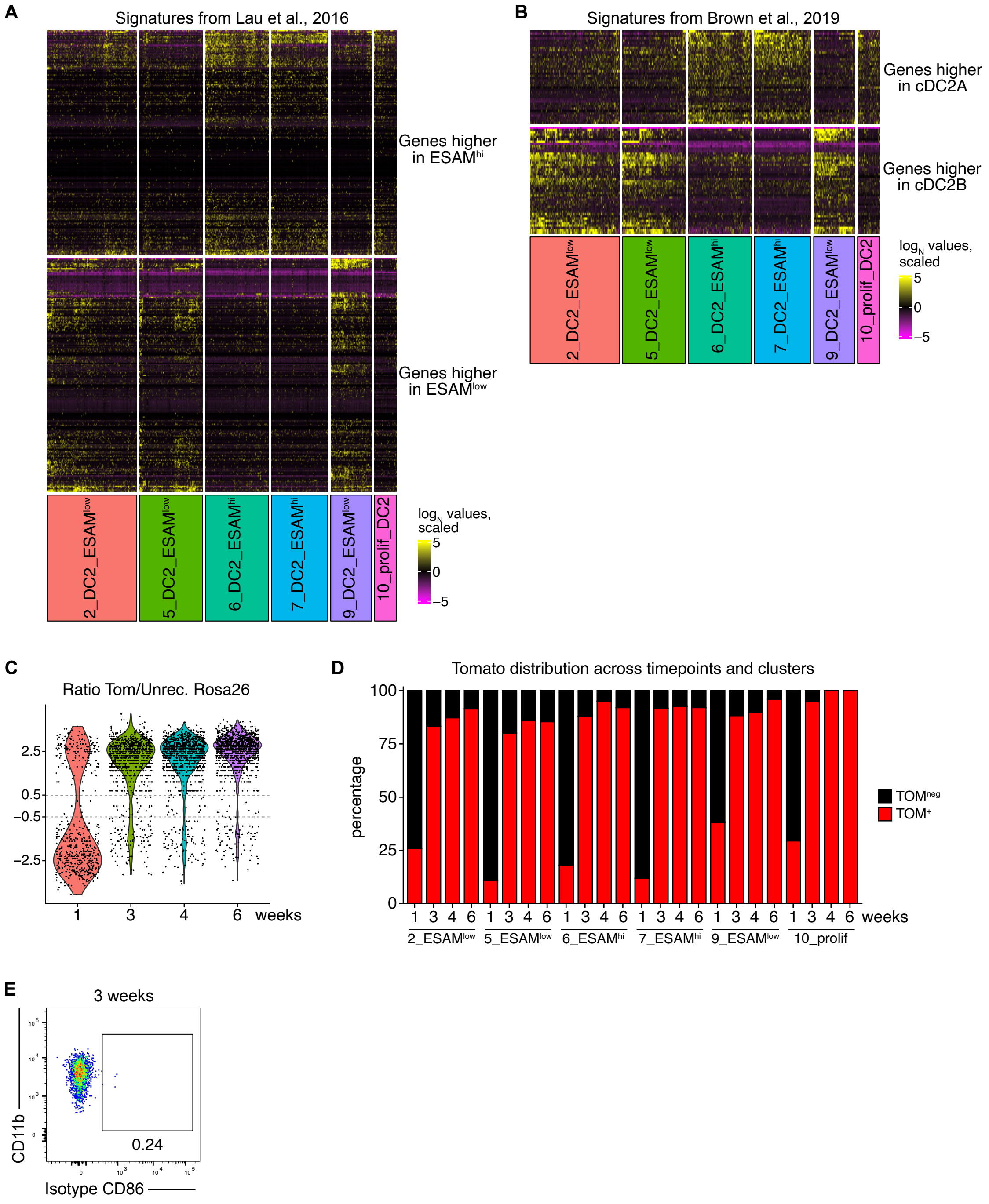

**Supplementary Figure 2:**

cDC2 and tDC were computationally isolated from a scRNA-seq data set of CD11c$^{+}$MHCII$^{+}$ splenocytes at 1, 3, 4 and 6 weeks of age. **(A)** Heatmap of signature genes for ESAM$^{hi}$ and ESAM$^{low}$ cDC2 identified by Lau et al., 2016 on cDC2 clusters. **(B)** Heatmap of signature genes for DC2A and DC2B identified by Brown et al., 2019. **(C, D)** All cDC2 clusters were grouped and the ratio of normalized TOMATO reads per cell to normalized reads per cell of the predicted transcript of the unrecombined ROSA locus was calculated. Cells with a ratio > 0.5 were identified as TOM+, whereas cells with a ratio <−0.5 were identified as TOM$^{neg}$. cDC2 with a ratio between -0.5 to 0.5 were classified as 'undefined'. **(D)** Frequency of TOM$^{+}$ and TOM$^{neg}$ cDC2 in the different cDC2 clusters, split by age. cDC2 with a ratio between -0.5 to 0.5 could not clearly be delineated as TOM$^{+}$ or TOM$^{neg}$ and are thus not shown on the quantification. **(E)** Fluorescent signal on ESAM$^{hi}$ cDC2A from 3-week-old mice stained with isotype antibody for anti-CD86 (related to Figure 2G, H).

### Supplementary Figure 3:

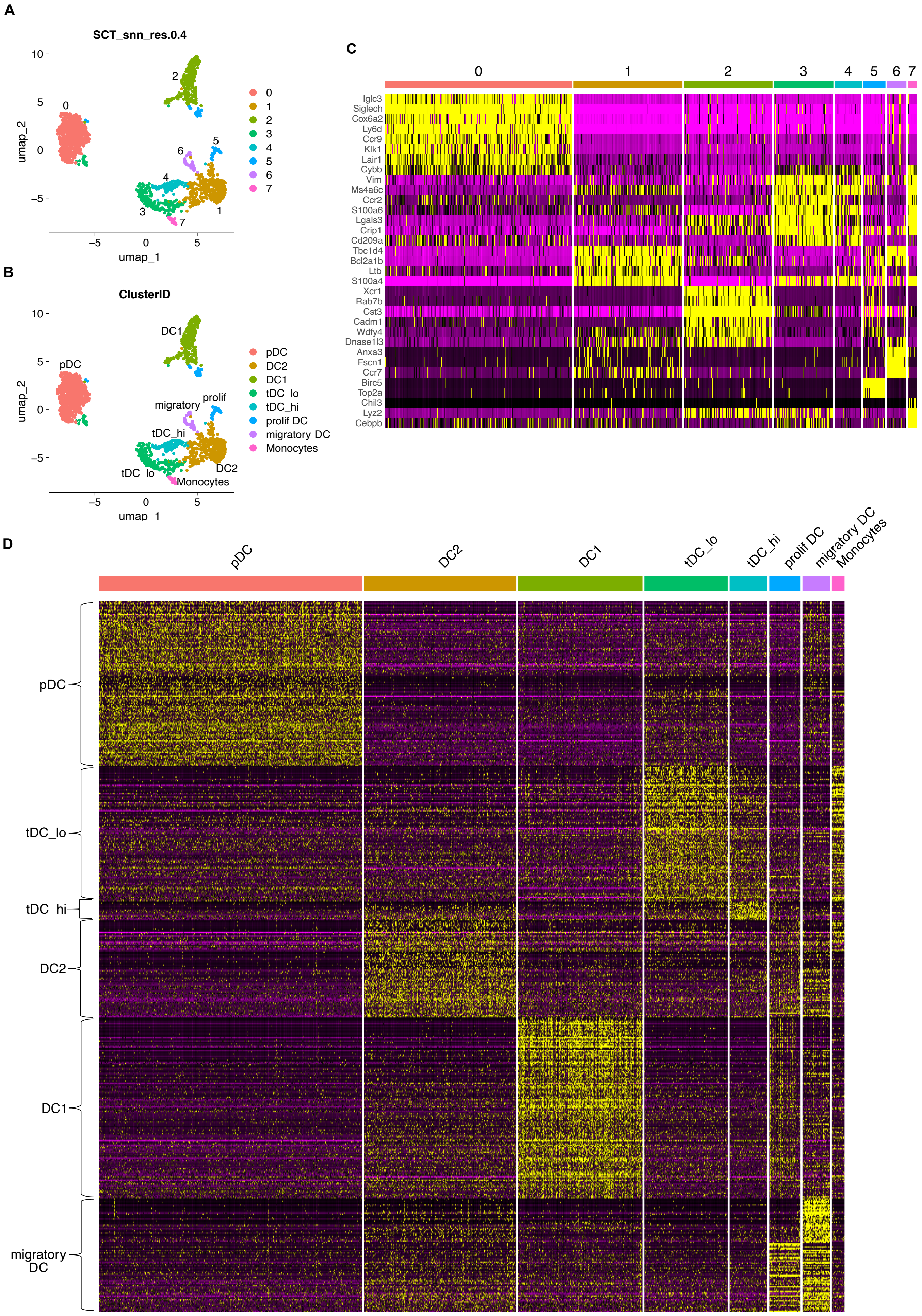

**Supplementary Figure 3:**

Cell type annotation for the scRNA-seq data from Sulczewski et al. show concordant cell types as defined in the original publication. **(A)** UMAP visualization of unsupervised clustering at the indicated resolution. **(B)** UMAP visualization grouped by cell type annotations. **(C, D)** Strategy for annotating cell types in **(B)**. **(C)** Heatmap showing genes used for cell type annotation by Sulczewski et al. in the clusters identified in **(A)**. **(D)** Heatmap showing differentially expressed genes between DC subtypes reported by Sulczewski et al.

### Supplementary Figure 4:

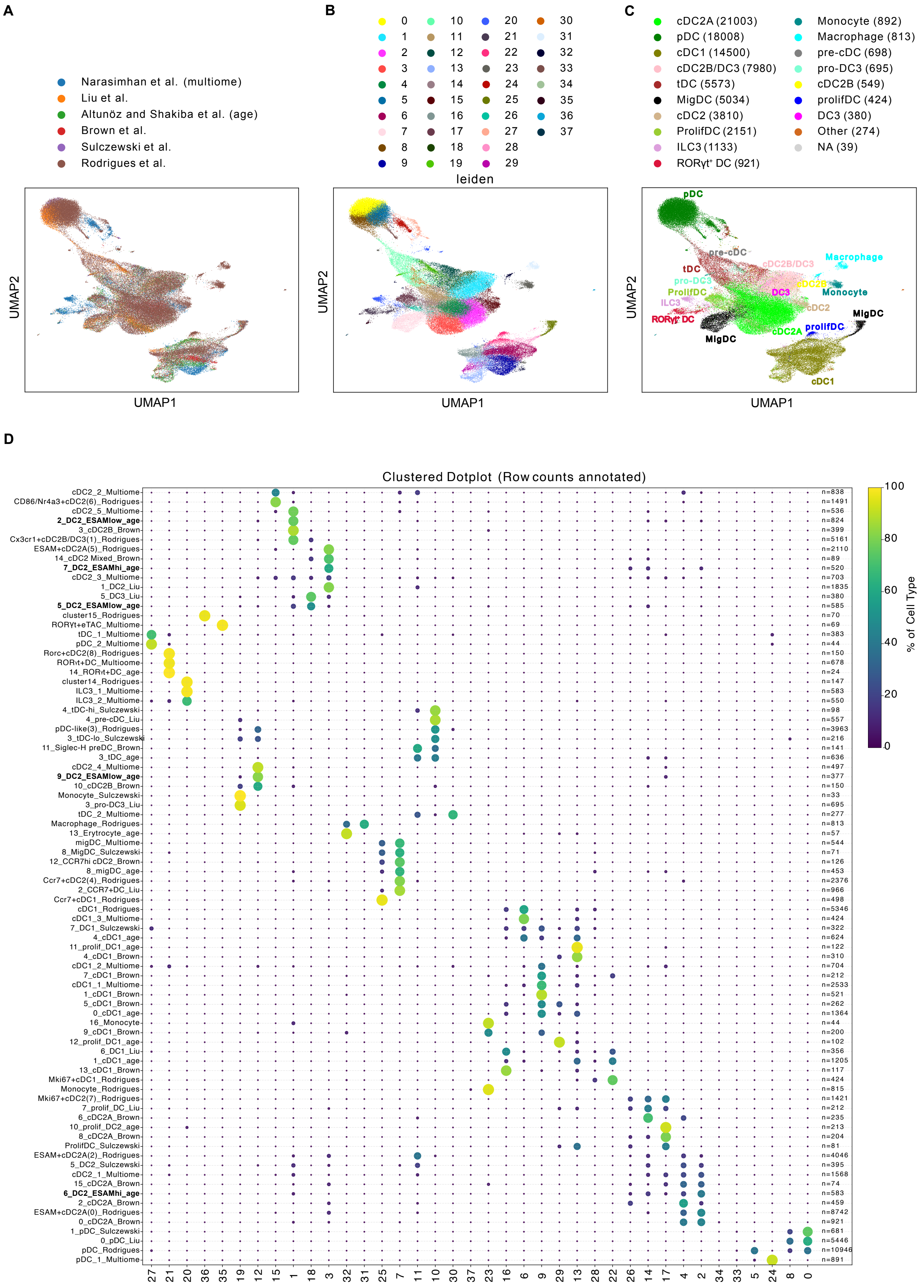

**Supplementary Figure 4:**

UMAP visualizations of the integrated data grouped by **(A)** dataset of origin, **(B)** leiden clusters, and **(C)** cell-type annotations based on the original publications (parentheses contain the number of cells per annotation). **(D)** Dot-plot summarizing the distribution of each annotated cell type from original datasets on the leiden clusters (sum of each row equals 100%). In addition, cell numbers of annotated cell types from original datasets are shown.

### Supplementary Figure 5:

**A**

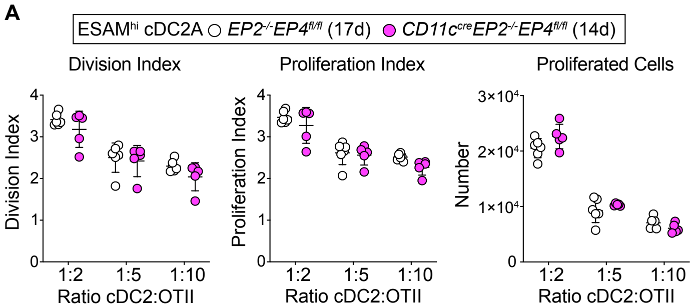


**B**

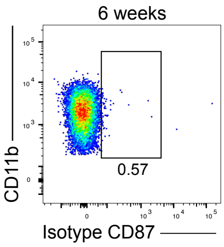

**Supplementary Figure 5:**

**(A)** Splenic ESAM$^{hi}$ cDC2A from mice of indicated genotypes were pulsed with $OVA_{323-339}$ and co-cultured with CTV-labelled OT-II cells under Th17 polarizing conditions at the indicated APC:T cell ratios for 3.5 days. The number of proliferated CTV$^{neg}$ cells, division and proliferation indices are shown. Data is representative of three independent experiments. Each dot represents one biological replicate. Horizontal bars represent mean, error bars represent SD. Statistical analysis was performed using two-tailed Welch's t tests with correction for multiple comparisons. *$p<0.05$, **$p<0.01$, ***$p<0.001$, ****$p<0.0001$. **(B)** Fluorescent signal on ESAM$^{hi}$ cDC2A from 6-week-old mice stained with isotype antibody for anti-CD87 (related to Figure 3G).

### Supplementary Figure 6:

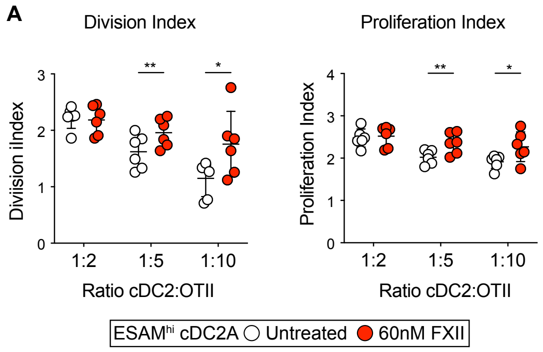

**Supplementary Figure 6:**

**(A)** Splenic ESAM$^{hi}$ cDC2A from 2-week-old mice were pulsed with $OVA_{323-339}$ and co-cultured with CTV-labelled OT-II cells under Th17 polarizing conditions at different DC:T cell ratios in the presence or absence of 60nM FXII for 3.5 days. Proliferation and division indices are shown. Data is representative of two independent experiments. Each dot represents one biological replicate. Statistical analysis was performed using two-tailed Welch's t tests with correction for multiple comparisons. *$p<0.05$, **$p<0.01$, ***$p<0.001$, ****$p<0.0001$.

### Supplementary Figure 7:

**A**

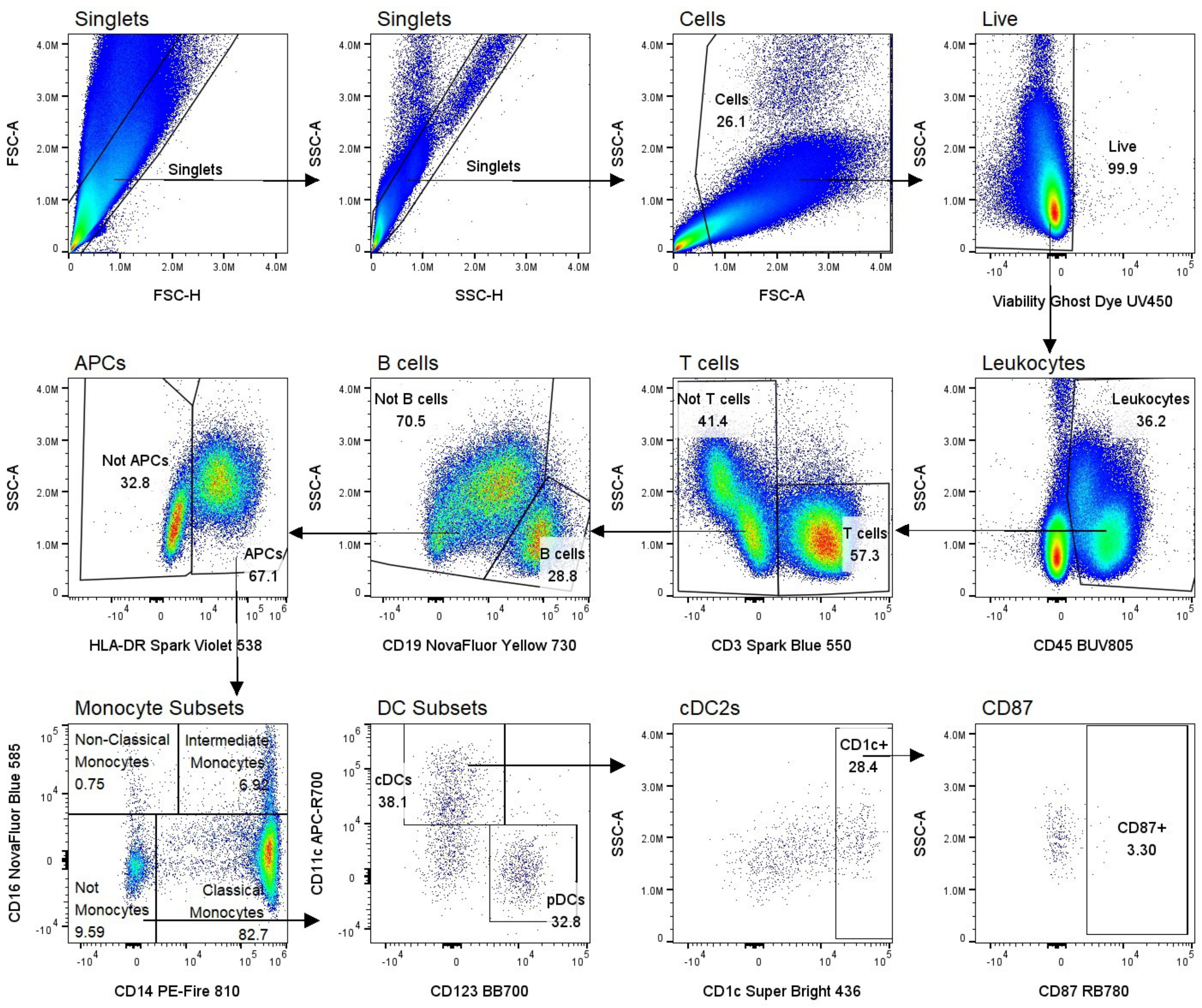

**Supplementary Figure 7:**

**(A)** Representative gating strategy to identify CD1c$^{+}$ cDCs in PBMCs and expression of CD87. First, doublets were removed by comparing FSC-A (forward-scatter-area) to FSC-H (forward-scatter-height), and then by SSC-A (side-scatter-area) to SSC-H (side-scatter-height). Debris was removed by gating out low SSC-A and FSC-A events, and dead cells were excluded. Leukocytes were identified by CD45. Live CD45$^{+}$ cells were then gated to CD3$^{neg}$CD19$^{neg}$HLA-DR$^{+}$ myeloid antigen-presenting cells (APCs) before removing monocyte subsets and other DC subsets to identify CD1c$^{+}$ cDCs as CD3$^{neg}$CD19$^{neg}$HLA-DR$^{+}$CD14$^{neg}$CD16$^{neg}$CD123$^{neg}$CD11c$^{+}$CD1c$^{+}$. CD87 expression was assessed within CD1c$^{+}$ cDCs.
